# Active removal of inhibitory components drives the flagellar Type III Secretion Specificity Switch

**DOI:** 10.64898/2026.03.09.710671

**Authors:** Fabienne F.V. Chevance, Dara Niketic, DingBang Wu, Carl Ty Mellor, David F. Blair, Sherwood R. Casjens, Miki Kinoshita, Keiichi Namba, Tohru Minamino, Kelly T. Hughes

**Affiliations:** School of Biological Sciences, The University of Utah, Salt Lake City, UT 84112, USA; Graduate School of Frontier Biosciences, The University of Osaka, Suita, Osaka 565-0871, Japan; JEOL YOKOGUSHI Research Alliance Laboratories, The University of Osaka, Suita, Osaka 565-0871, Japan

## Abstract

Type III secretion (T3S) systems assemble bacterial nanomachines, including the flagellum and virulence-associated injectisomes, by exporting distinct classes of substrates in a defined temporal order. In both systems, completion of an early assembly intermediate triggers an irreversible switch from early to late substrate secretion. In the flagellar system, this switch is controlled by the secreted molecular ruler FliK acting on the core T3S component FlhB, but the molecular mechanism governing this transition has remained unclear. Here we show that removal of two components, Fluke and the cleaved C-terminal domain of FlhB (FlhB_CCD_), locks the secretion apparatus in a constitutive late secretion state. In these mutants, secretion specificity no longer requires completion of the hook-basal body or the FliK ruler, indicating that Fluke and FlhB_CCD_ function to maintain the apparatus in early secretion mode. Consistent with this model, synchronized flagellar gene expression experiments reveal that FlhB_CCD_ is retained during early assembly and is lost coincident with hook-basal body completion and activation of σ^28^-dependent late gene expression of flagellin and chemosensory genes. Structural modeling of the FliK C-terminal switch domain and FlhB_CCD_ supports a mechanism in which secretion of FliK promotes destabilization and ejection of FlhB_CCD_ from the secretion apparatus. Disruption of a folded region within FliK switch domain uncouples secretion from switching, indicating that the timing of FliK unfolding during secretion is critical for activation of the specificity switch. These findings show that secretion specificity switching is driven by FliK-dependent removal of inhibitory components, rather than passive sensing of assembly completion.

**Significance Statement:** Type III secretion (T3S) systems build complex bacterial nanomachines, including the flagellum and virulence-associated injectisomes, by exporting distinct classes of substrates in a defined temporal order. How these systems switch secretion specificity during assembly has remained a long-standing question. We demonstrate that the flagellar T3S specificity switch requires removal of two inhibitory components that actively maintain the apparatus in an early secretion state. Their FliK-dependent ejection irreversibly triggers the transition to late substrate export, revealing that secretion switching is controlled by active inhibitory regulation rather than passive sensing of assembly completion. These results define a molecular mechanism for hierarchical control in bacterial secretion systems and provide insights likely relevant to other T3S nanomachines.

## Introduction

Many bacteria are motile, using external flagellar organelles to swim in liquid environments and swarm across hydrated surfaces (5). Flagella are large macromolecular assemblies that extend ∼10 microns from the cell surface and are constructed by the secretion and ordered self-assembly at the distal tip of the growing structure (37). In enteric bacteria, fourteen different proteins are exported from the cytoplasm through the flagellar type III secretion (T3S) system, a specialized secretion apparatus that coordinates assembly of the flagellum in a defined temporal order. The flagellar T3S system is a rare example of a secretion system that undergoes an irreversible switch in secretion specificity, transitioning from early substrates required for rod-hook assembly to late substrates that form the filament and associated structures.

The secretion-specificity switch occurs upon completion of the hook-basal body (HBB), a conserved assembly intermediate that anchors the flagellum in the cell envelope (**Figure 1**). During early assembly, rod and hook subunits are secreted through the T3S apparatus and polymerize beneath the transient cap structures at the distal end of the growing organelle (reviewed in (8)). Upon hook completion, secretion switches to late substrates, including hook-filament junction proteins, the filament cap, and filament subunits (9; 12). All structural components of the rod-hook-filament axial structure are transported through a narrow central channel and assembly exclusively at the distal tip.

**Figure 1.**
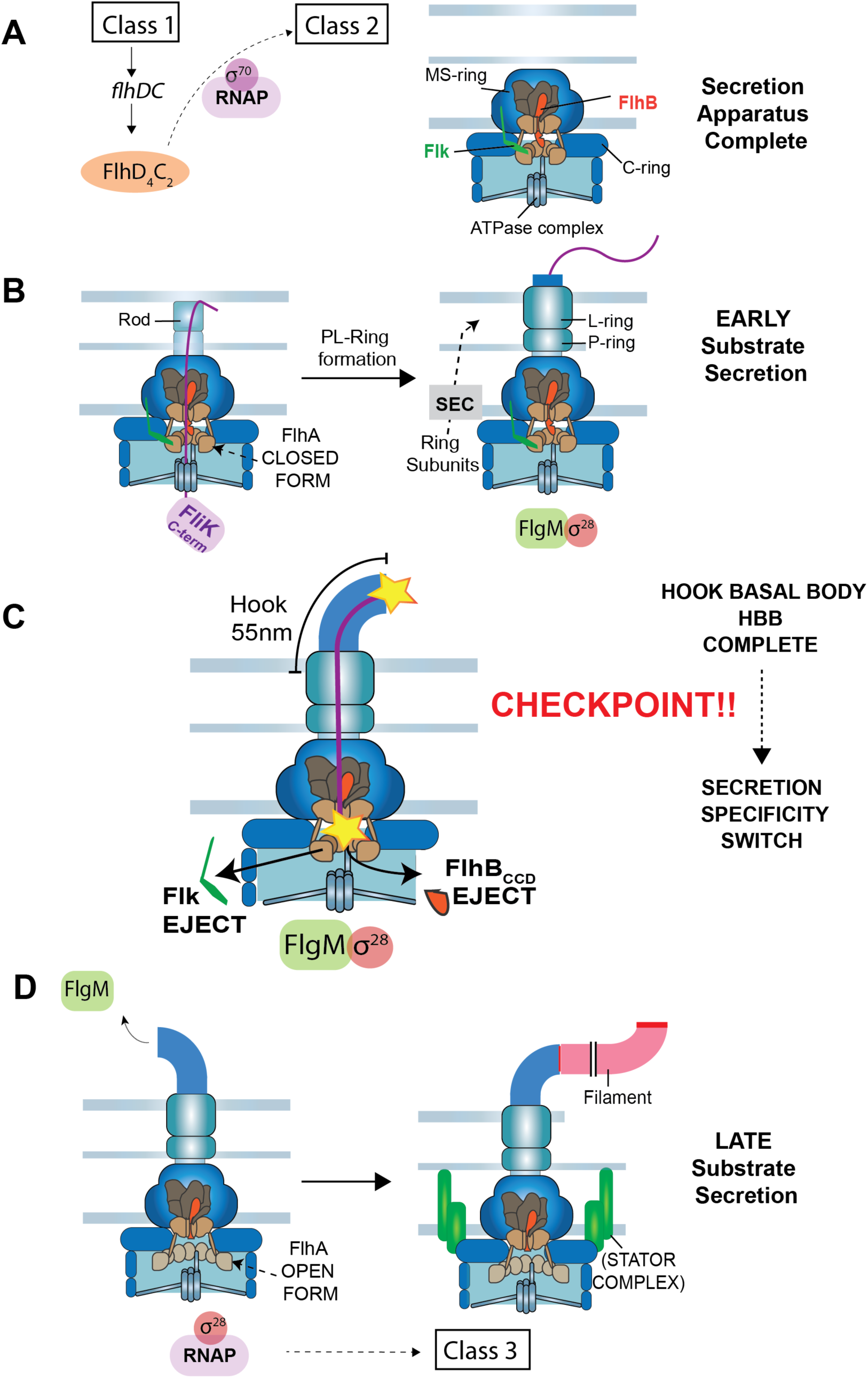
Flagellar biogenesis in *Salmonella* and coupled gene regulatory mechanisms. **A.** Flagellum assembly initiates with expression of the class 1 *flhDC* master operon. The *flhDC* genes encode a transcriptional activation complex that directs RNA polymerase to transcribe flagellar class 2 promoters. Gene expressed from class 2 flagellar promoters encode the proteins necessary for the formation of the hook-basal body (HBB) intermediate structure. Assembly begins with the formation of the secretion apparatus core in the inner membrane. The core is composed of FliP_5_-FliQ_4_-FliR (shown in dark brown), which is associated with a single C-terminal domain of FlhB (colored red). Beneath the T3S core is a nonameric ring of FlhA (colored tan). The N-terminal region of FlhA (FlhA_N_, residues 1 – 327) is composed of predicted transmembrane segments. The FlhA C-terminal ring (residues 363 – 692) extends below, connected to FlhA_N_ by a flexible linker region (residues 328 – 361). The T3S core is contained within the MS-ring (FliF) in the inner membrane. Beneath the MS-ring is the flagellar rotor also called the C-ring composed of FliG, FliM and FliN proteins. At the cytoplasmic base of the structure is the FliH-FliI-FliJ ATPase complex, which delivers substrates to FlhA for secretion. **B.** The flagellar rod structure extends from the T3S core through the cell wall to the outer membrane. The FliK molecular ruler is intermittently secreted during rod-hook formation taking temporal measurements of the growing structure. Outer membrane penetration requires the formation of the PL-ring around the distal rod to form the outer membrane pore. The proteins required for PL-ring formation are secreted into the periplasm by the general Sec secretion system. The gene encoding the σ^28^ transcription factor required for class 3 flagellar gene transcription is expressed along with HBB genes from a flagellar class 2 promoter, but it is held inactive by the anti-σ^28^ factor FlgM. **C.** Following PL-ring completion a flexible hook structure extends from the cell surface. Hook subunits are secreted and their polymerization terminates when a secreted FliK molecule measure a minimal length (between the yellow stars in figure) that places the C-terminus of FliK (FliK_C_) in the vicinity of the T3S core. FliK_C_ catalyzes the removal of the Fluke and FlhB_CCD_ “locks” to flip the secretion-specificity switch from early to late substrate specificity. Prior to hook completion, secretion of FliK occurs at a rate that is too fast to allow a productive interaction between FliK_C_ and the core to flip the secretion-specificity switch. Removal of the Fluke and FlhB_CCD_ “locks” results in a conformational change in FlhA_C_ from a closed to an open conformation that allows delivery of late secretion substrates by their bound T3S chaperones. **D.** FlgM is a late secretion substrate that is secreted upon HBB completion releasing σ^28^ to direct RNA polymerase to flagellar class 3 promoters. Assembly of the MotAB stator complexes in the inner membrane and polymerization of the long external filament from the hook completes the biogenesis of the *Salmonella* flagellum.

Completion of the HBB is sensed by the secreted molecular ruler, FliK (18), which determines hook length and triggers the secretion-specificity switch. Multiple FliK molecules are secreted during hook assembly (39), and the final hook length corresponds to the physical length of FliK (51). Productive switching requires interaction the C-terminal switch domain of FliK (FliKC) and the core T3S component FlhB, which resides at the cytoplasmic base of the secretion apparatus (33). Current models propose that when a secreting FliK molecule pauses upon reaching the distal tip of a fully elongated hook, FliK_C_ is positioned near FlhB long enough to trigger switching (67). However, how this interaction converts hook-length sensing to an irreversible change in secretion specificity remains unresolved.

FlhB is a 384-residue protein that undergoes autocleavage between N269 and P270, generating a cleaved C-terminal domain (FlhB_CCD_) that is essential for secretion switching (41). Mutations that disrupt FliK function or prevent FlhB autocleavage result in a polyhook phenotype, reflecting a failure to transition from early to late secretion (41; 45). Notably, mutations within FlhB_CCD_ can partially bypass the requirement for FliK, allowing inefficient late substrate secretion even in its absence (22; 35). These observations led to the proposal that FliK_C_ promotes switching by inducing a conformational change in FlhB_CCD_ (42).

In addition to FlhB, secretion switching is accompanied by a conformational change in the FlhA component of the secretion apparatus. FlhA forms a cytoplasmic nonameric ring that serves as a docking platform for late secretion substrates delivered by dedicated T3S secretion-chaperones (**Figure 1**) (4; 19; 29; 48). Prior to switching, FlhA adopts a closed conformation that occludes the secretion-chaperone binding site. Following switching, FlhA transitions to an open state that permits targeted delivery of late substrates (24; 32; 56). This conformational change is dependent on FliK, linking hook-length sensing to late substrate secretion.

Secretion switching is also tightly coordinated with transcriptional control of flagellar gene expression. The alternative transcription factor σ^28^ directs transcription of filament and chemotaxis genes but is inhibited prior to HBB completion by the anti-sigma factor FlgM (44). FlgM is itself a late secretion substrate and is secreted only after the specificity switch, thereby coupling filament assembly and chemotaxis gene expression to completion of the HBB structure (23). The protein, Flk (also known as RflH; hereafter referred to as Fluke) has been implicated in preventing premature FlgM secretion, suggesting that active mechanisms exist to maintain the early secretion state prior to HBB completion (2; 34).

Here we show that Fluke and FlhB_CCD_ function as inhibitory components that actively maintain the flagellar T3S apparatus in early secretion mode. We demonstrate that FliK-dependent removal of both components irreversibly triggers the secretion-specificity switch upon HBB completion. Furthermore, we show that switching requires both a secretion pause of FliK at full hook length and the unfolding of a discrete domain within FliK_C_, providing the time necessary for productive interaction with FlhB_CCD_. Together, our findings define a molecular mechanism by which assembly-state sensing is converted into irreversible secretion switching during flagellar biogenesis.

## Results

### Fluke inhibits FliK-dependent T3S-specificity switching prior to HBB completion

In addition to FlhB autocleavage and interaction between FliK_C_ and FlhB_CCD_ causing the secretion-specificity switch to flip (31; 41), we previously identified a third protein Fluke as an inhibitor of late secretion but the mechanism of action was not known (27). Fluke is a 333 amino acid cytoplasmic-facing membrane anchored protein (2). The absence of Fluke in a *flk*^-^ strain results in a premature switch in T3S specificity to late secretion mode without requiring HBB completion as is demonstrated by the secretion of late substrate FlgM into the periplasm in a Δ*flgHI* PL-ring mutant (“PL-ring^-^“ hereafter) (3). Despite this, strains deleted only for the *flk* gene exhibit the same motility phenotype on soft agar swim plates as *flk*^+^ cells (**Supplemental Figure 1**). Since wild-type *Salmonella* have an average of four flagella per cell, but only one functional flagellum is required for motility (17; 54), these two seemingly conflicting findings might be resolved if Fluke inhibition is incomplete, and a subset of basal body structures undergo the change in specificity and assemble full external flagella in the absence of Fluke. We examined this by measuring switch flipping as indicated by early and late flagellar gene expression with and without the PL-ring (**Figure 1**) as follows: Cells expressing early (Class 2) and late (Class 3, σ²⁸-dependent) secretion genes were quantified using fluorescent transcriptional reporters. Expression of early secretion substrates was monitored with a YFP fusion to the Class 2 *fliF* promoter (P*_fliF_*-*yfp*; Fig. 1A), while late secretion gene expression was monitored with a CFP fusion to the Class III *fliC* promoter (P*_fliC_*-*cfp*), which is activated following secretion of the anti-σ²⁸ factor FlgM (**Figure 2A**). Because YFP expression precedes CFP expression, CFP-positive cells are expected to also express YFP. Consistent with this, in wild-type cells >99% of P*_fliF_*-*yfp* positive cells also expressed the Class 3 reporter (**Figure 2B**), indicating that essentially all Class 3 expressing cells had undergone the T3SS specificity switch.

**Figure 2.**
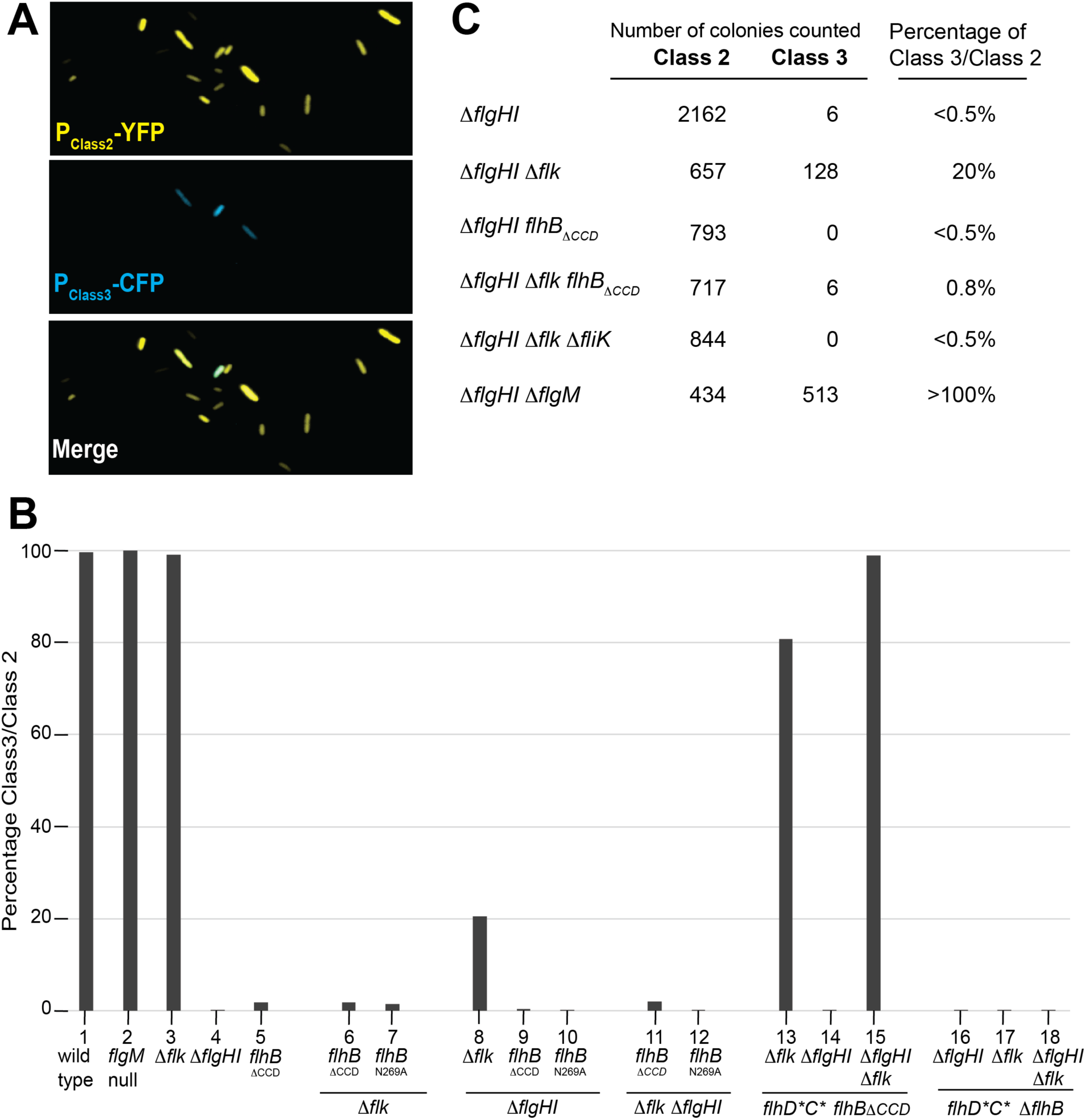
Class 3 promoter activity relative to class 2 promoter activity in individual bacterial cells, in different genetic backgrounds. **A.** An example of microscopic images of individual cells expressing class 2 genes (YFP) and class 3 genes (CFP) in TH28053 (*attB*::[Cm P*_fliF_*-*yfp*] Δ*galK*::[Ap P*_fliC_*-*cfp*] Δ*flgHI*958 Δ*flk*-7755 *fljB^enx^ vh2*). **B.** Percentage of class 3 promoter activity relative to class 2 promoter activity in individual bacterial cells from different genetic backgrounds. The number of cells expressing class 3 activity was counted relative the number of cells expressing class 2 activity in strains of various genetic backgrounds and shown in the following lanes: 1=TH28049 (wild type); 2=TH27343 (*flgM* null); 3=TH28052 (Δ*flk)* ; 4=TH28050 (Δ*flgHI*); 5=TH28051 (*flhB*_ΔCCD_); 6=TH28054 (Δ*flk flhB*_ΔCCD_); 7=TH29314 (Δ*flk flhB*_N269A_); 8=TH28053 (Δ*flk* Δ*flgHI*); 9=TH28055 (Δ*flgHI flhB*_ΔCCD_); 10=TH29315 (Δ*flgHI flhB*_N269A_) ; 11=TH28056 (Δ*flk* Δ*flgHI flhB*_ΔCCD_); 12=TH29316 (Δ*flk* Δ*flgHI flhB*_N269A_); 13=TH29317 (*flhD*\*C* Δ*flk flhB*_ΔCCD_); 14=TH29318 (*flhD*\*C* Δ*flgHI flhB*_ΔCCD_); 15=TH29319 (*flhD*\*C* Δ*flk* Δ*flgHI flhB*_ΔCCD_); 16=TH29357 (*flhD*\*C* Δ*flk* Δ*flhB*); 17=TH29358 (*flhD*\*C* Δ*flgHI* Δ*flhB*); 18=TH29359 ((*flhD*\*C* Δ*flk* Δ*flgHI* Δ*flhB*). All strains contained *attB*::[Cm P*_fliF_*-yfp] Δ*galK*::[Ap-P*_fliC_*-*cfp*] and their complete genotypes are listed in Table S1. **C.** Percentage of class 3 promoter activity relative to class 2 promoter activity in individual bacterial cells with no PL-rings (Δ*flgHI*). The number of cells expressing class 3 activity was counted relative the number of cells expressing class 2 activity in ring mutant backgrounds cells (TH27356 (Δ*flgHI*), TH27359 (Δ*flk* Δ*flgHI*), TH27477 (Δ*flgHI flhB*_ΔCCD_), TH27476 (Δ*flgHI* Δ*flk flhB*_ΔCCD_), TH27478 (Δ*flgHI* Δ*flk*, Δ*fliK*), TH27451 (Δ*flgHI* Δ*flgM*). These strains contained *attB*::[Km^R^-P*_fliF_*-yfp] Δ*galK*::[Ap^R^-P*_fliC_*-*cfp*] and their complete genotypes are listed in Table S1.

In a *flk*^+^ PL-ring^-^ strain where the HBB is incomplete and secreted proteins remain in the periplasm (above), only 0.3% of the cells that expressed YFP (Class 2) also expressed CFP (Class 3) indicating that nearly all cells expressing Class 2 genes had *not* undergone the T3S specificity switch as a result of ongoing FlgM-dependent inhibition of class 3 gene expression (**Figure 2B, 2C**). On the other hand, in a *flk*^-^ PL-ring^-^ strain 20% of the cells that expressed YFP also expressed CFP indicating that a fifth of the cells the T3S-specificity switch was flipped to late mode secretion (**Figure 2B, 2C**). Class 3 CFP expression in the Δ*flk* PL-ring^-^ background remained dependent on the presence of the FliK ruler and a functional FlhB (**Figure 2C**). These findings show that Fluke prevents growing basal body structures from undergoing a premature FliK-dependent secretion specificity change before completion of the HBB. However, in Fluke’s absence release from this inhibition is incomplete, so something else continues to exert partial inhibition of switching in Fluke’s absence.

### Fluke or FlhB_CCD_ prevents late substrate secretion independently from one another

The absence of Fluke removes one source of inhibition of late secretion. Therefore, selection in the absence of Fluke for mutations that allow Class 3 gene expression (a consequence of late secretion) should identify other inhibitors of the switch to late secretion mode. To search for such mutations, we used the system we developed that utilizes β-lactamase (Bla) lacking its Sec secretion signal but fused to a flagellar gene secretion signal as a reporter for T3S-mediated secretion (10). When the rod structure is missing in a Δ*flgB-L* strain, secretion is directed into the periplasm (Δ*flgB-L,* a.k.a. “Rod-hook^−^”, is deleted for all the genes encoding rod-hook proteins but leaves the secretion apparatus intact; unlike PL-ring^−^ above, the Rod-hook^−^ deletion does not allow partial late secretion in the absence of Fluke (above)). In this system, fusion to a flagellar gene’s secretion signal allows secretion of Bla into the periplasm where it folds into an active conformation and confers resistance to ampicillin (Ap^R^). Thus, a *flgM-bla* fusion that has the FlgM *late* gene secretion signal provides a positive Ap^R^ selection for cells performing late secretion. In addition, to maximize secretion of FlgM-Bla and make selection more efficient (i) the selection strain was deleted for all other late secretion substrates (Δ*fliB-T fljB^enx^ vh2,* a.k.a “Late substrate^-^“) that might compete with it for secretion, (ii) a mutation was included in the σ^28^ structural gene *fliA* (*fliA5225*(H14D) a.k.a “σ^28^1↑”) which results in a 2-fold increase in the σ^28^ levels (7), and σ^28^ also facilitates FlgM secretion in addition to its role as a late transcription activator (3), and (iii) protease-resistant alleles of the master regulators that result in ∼4-fold more HBB structures per cell (*flhD8070*(L22H), *flhC8092*(Q29P), a.k.a. “*flhD*\**C*\*”) were included (50; 55).

This final *flk*^-^ selection strain (TH25226: *flk*^-^ FlgM-Bla Rod-hook^−^ *flhD*\**C*\* σ^28^↑ Late substrate^-^) was, as expected, in early secretion mode and thus Ap^S^. When plated on ampicillin selection medium, Ap^R^ mutants that secrete FlgM-Bla into the periplasm arose. Among the mutants obtained, two classes were unrelated to the switch: (i) changes in SecY that allowed FlgM-Bla which has no Sec secretion signal to be secreted through the Sec pathway into the periplasm and (ii) deletions that fused the Sec secretion signal from the FlgA protein to FlgM-Bla, which also resulted in Sec-dependent Bla secretion (**Figure 3A**). For the third Ap^R^ mutant class, introduction of a TS3 secretion apparatus null allele (Δ*fliP*) resulted in an Ap^S^ phenotype, indicating that this apparatus was required for FlgM-Bla secretion. Thus, this class of mutants performs late secretion without completing HBB assembly (since no rod-hook proteins are made). Ten of the latter Ap^R^ mutations were mapped on the *Salmonella* chromosome, and their DNA sequences were analyzed, which revealed that all ten have an altered FlhB_CCD_ (**Figure 3B**). To our surprise one of these mutations is a stop codon in *flhB* at residue 258, which causes the loss of the entire FlhB_CCD_. To confirm this result, a complete deletion (ΔAA270 - 459) of FlhB_CCD_ was introduced into the original selection TH25226 strain, and it resulted in an Ap^R^ phenotype (**Figure 3C, 3D**). This suggests that it may not be a conformational change in FlhB_CCD_ that causes flipping of the specificity switch as was previously thought, but its physical removal may be required to flip the switch. This would explain why FlhB undergoes autocleavage (which is required for normal late secretion (41)), since a covalent linkage to the rest of FlhB would deny the physical removal of FlhB_CCD_.

**Figure 3.**
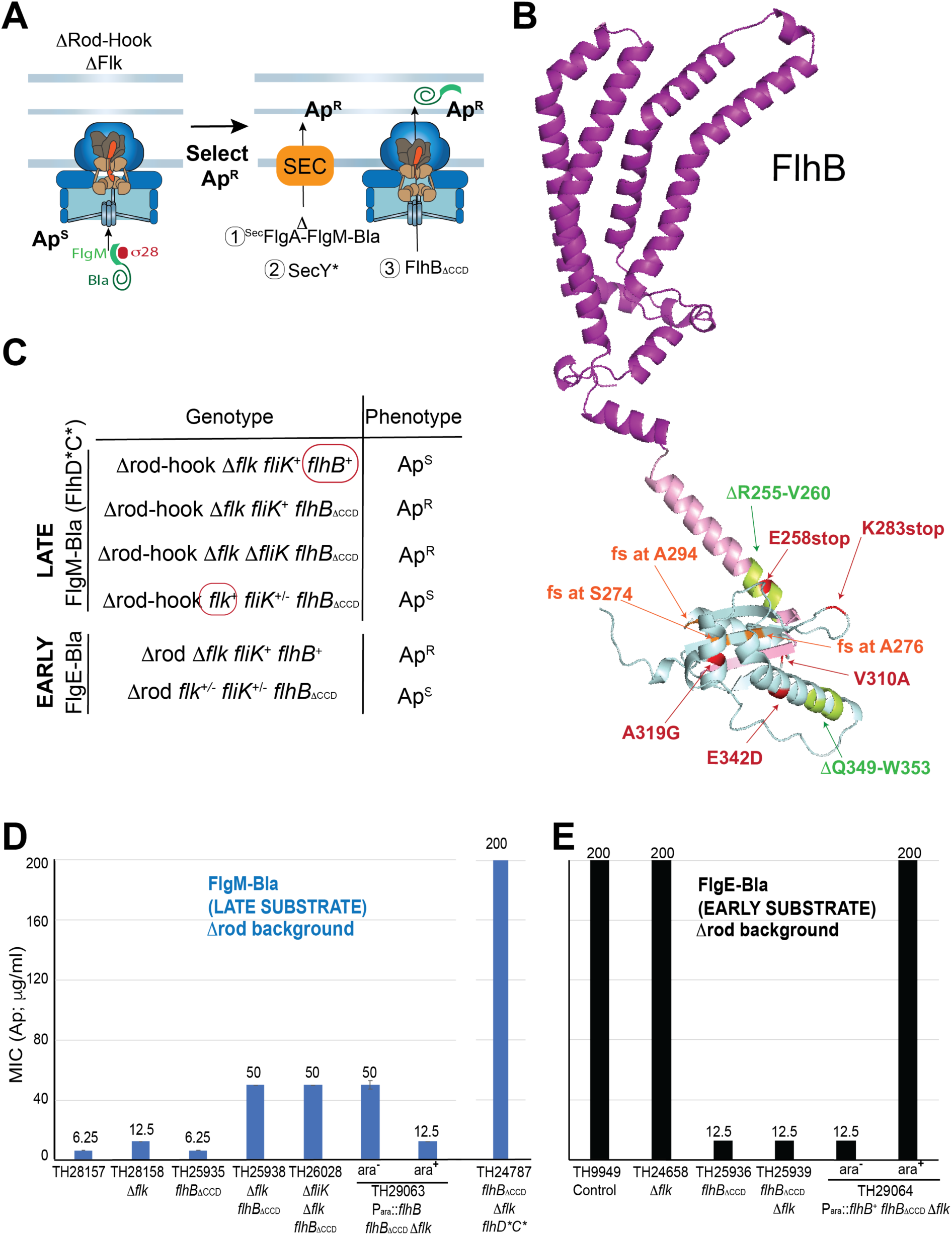
Selection for FlgM-bla secretion results in FlhB C-terminal domain mutants including null alleles lacking the C-terminal domain. **A.** Selection strategy used to obtain mutants permitting the secretion of late flagellar substrates. In an HBB mutant strain, flagellar late-secretion substrates are not secreted. Thus, when the late substrate FlgM is fused to mature β-lactamase (Bla), the strain is ampicillin sensitive. In a strain lacking both the HBB and the Flk protein, it is possible to obtain Ap^R^ mutants. **C.** Mutants in the flagellar type 3 secretion apparatus that allow secretion of flagellar late-secretion substrates located in the C-terminal domain of FlhB. A large portion of the mutants obtained in the C-terminal domain of FlhB are stop codons or frame-shift mutations resulting in premature stop codons (mutations illustrated in green). **D.** The secretion of FlgM-bla (late flagellar substrate), using minimum inhibitory assays of ampicillin (10) in various mutant backgrounds. The secretion of FlgM-bla is increased 4-fold in the *flhD*C\** Δ*flk flhB*_ΔCCD_ mutant background (far right strainTH24787). **E.** Effect of the deletion of the C-terminal domain of FlhB (FlhB_ΔCCD_) on the secretion of early (FlgE-Bla).

The selection strain lacks all rod-hook structural genes. Thus, we also tested the requirement for the FliK ruler since there was no structure to measure. Deletion of *fliK* had no effect on FlgM-Bla secretion (**Figure 3C, 3D**). However, when a *flk*^+^ allele and P*_fliC_*-*cfp* were introduced into the above selection strain that was also deleted for FlhB_CCD_ the cells did not express CFP (**Figure 2B**) and became Ap^S^ (**Figure 3C, 3D**) indicating that either FlhB_CCD_ or Fluke is sufficient to prevent the switch to late secretion (**Figure 2B**). The fact that FliK is not required for late secretion under these conditions is consistent with a model in which FliK’s role in flipping the switch is to inactivate or remove FlhB_CCD_, and Fluke inhibition does not require FliK or FlhB_CCD_ (see possible FlhA_C_ interaction below).

We note that HBB over-expression was required for FlgM-Bla secretion in the Δ*flk* Δ*flhB_CCD_* strain since replacing the hyperactive *flhD*C*\* alleles (above) with wild-type alleles resulted in a significant reduction of ampicillin resistance and class 3 P*_fliC_*-*cfp* gene expression (**Figure 2B, 3D**). This would be explained if FlhB normally assembles as an intact protein prior to autocleavage, and inefficient assembly of FlhB without its CCD is overcome by overexpression in the presence of more HBBs.

### Loss of FlhB_CCD_ results in a T3S structure locked in late secretion mode

Early substrates have been reported to be targeted for secretion by interacting with a hydrophobic pocket in FlhB_CCD_ (6; 47), so the loss of FlhB_CCD_ should disallow early secretion and perhaps lock the switch in late secretion mode (above). We tested this by introducing an *flhB* gene deleted for FlhB_CCD_ coding sequence (*flhB_ΔCCD_*) into a strain expressing a Bla fusion to the *early* secretion substrate hook protein (FlgE) (again lacking the Bla Sec secretion signal). Strains defective in rod assembly (Δ*flgBC*) secrete the FlgE-Bla fusion into the periplasm and are Ap^R^ (**Figure 3E**). The *flhB_ΔCCD_* allele caused an Ap^S^ phenotype, indicating that FlgE-Bla is not secreted in the absence of the FlhB_CCD_ (**Figure 3C and E**). The presence or absence of Fluke did not affect this result. This shows that the locked-in-late-secretion mode does not secrete early substrates, and the lack of FlhB_CCD_ results in a T3S structure locked in late secretion mode due to the lack of an early substrate docking site.

### Prevention of premature flipping of the specificity switch at the rod to hook assembly transition

The transition from internal rod completion to external hook assembly is a critical step in flagellum assembly. The transition from rod completion to hook polymerization requires Sec-dependent secretion of proteins that build the PL-ring. Formation of the PL-ring around the distal rod simultaneously forms an outer membrane pore and dislodges the rod cap protein (9; 12). The Fluke and FlhE proteins are also required for the efficient transition from rod-to-hook polymerization (3; 19). T3S-specificity switching in the absence of Fluke occurs in PL-ring^-^ strains **but** not when steps of HBB formation are blocked (above) (27). The loss of FlhE results in periplasmic flagella (21). This suggests that the role of FlhE is to maintain the rod cap prior to PL-ring formation thus preventing premature hook and filament protein secretion and polymerization prior to outer membrane penetration. In addition, overexpression of the hook structural gene *flgE* suppresses motility defects in PL-ring mutants and electron micrographs of isolated flagellar structures from a P-ring mutant with hook (FlgE) overexpression showed a structure lacking the PL-ring with a normal appearing rod-hook-filament transition (43). Finally, the rod’s tip reaching the outer membrane does not prevent the continuous secretion of early substrates into the periplasm (11; 36). These observations taken together suggest that, since rod completion and the transition to hook assembly are not strictly coupled, mechanisms are in place to prevent switching before PL-ring assembly and hook formation. The data in **Figure 2** imply that Fluke also plays a role in the rod-to-hook transition step in flagellar formation in that it helps prevent premature secretion of late substrates when the PL-ring is missing.

As described above, FlhA is a membrane protein whose cytoplasmic domain FlhA_C_ binds late substrate-chaperone complexes and presents the substates for secretion (29; 40; 58); however, in order to carry out this function, FlhA_C_ undergoes a FliK dependent conformational change to an “open” form that exposes the late substrate T3S chaperone docking site (24; 32; 56). Since Fluke is a cytoplasmic protein with a C-terminal membrane anchor, it is possible that it acts on FlhA_C_ to prevent the conformational change in FlhA that occurs upon hook completion (24). We tested for a possible Fluke-FlhA_C_ interaction by bacterial two-hybrid assay and obtained a positive interaction result that supports this idea (**Supplemental Figure 2**). In addition, FliK also exhibited a positive interaction with FlhA_C_ in the two-hybrid assay. This is consistent with the idea that part of Fluke’s function is to prevent FlhA from binding late secretion substrates prematurely. Currently, it is thought that FliK flips the switch not only by catalyzing removal of FlhB_CCD_ but changing the conformation of FlhA to the “open” late substrate secretion mode. It seems likely that FlhB_CCD_ holds FlhA in the “closed” early secretion mode and its removal allows FlhA to change to the open conformation. A more detailed characterization of Fluke-FlhA and FliK-FlhA interactions await future studies.

If FliK secretion is slowed in cells lacking the PL-ring, it might increase the probability that FliK_C_ flips the specificity switch prematurely. FliK is being continuously secreted during rod and hook assembly, and it seemed possible that FliK secretion into the periplasm in a PL-ring mutant (with no outer membrane pore) might be slowed when the rod-tip abuts the outer membrane. We tested this by measuring levels of FliK-Bla secreted into the periplasm in mutants defective in each step in assembly from rod initiation to filament polymerization (**Figure 4**). Using the minimal inhibitory concentration (MIC) of ampicillin (Ap) as an indicator of secreted FliK-Bla levels, a 2-fold reduction in periplasmic FliK-Bla activity was observed in strains defective in PL-ring assembly as compared to mutants defective in rod or hook assembly. Mutants defective in steps after PL-ring assembly showed an additional 2-fold reduction. This is the same level of FliK-Bla secreted into the periplasm observed in wild-type cells. We conclude that the low level of Ap^R^ in mutants defective in steps after PL-ring assembly (*flgK*^-^) likely reflects the low amount of FliK-Bla secreted during rod assembly prior to outer membrane penetration by the PL-ring.

**Figure 4.**
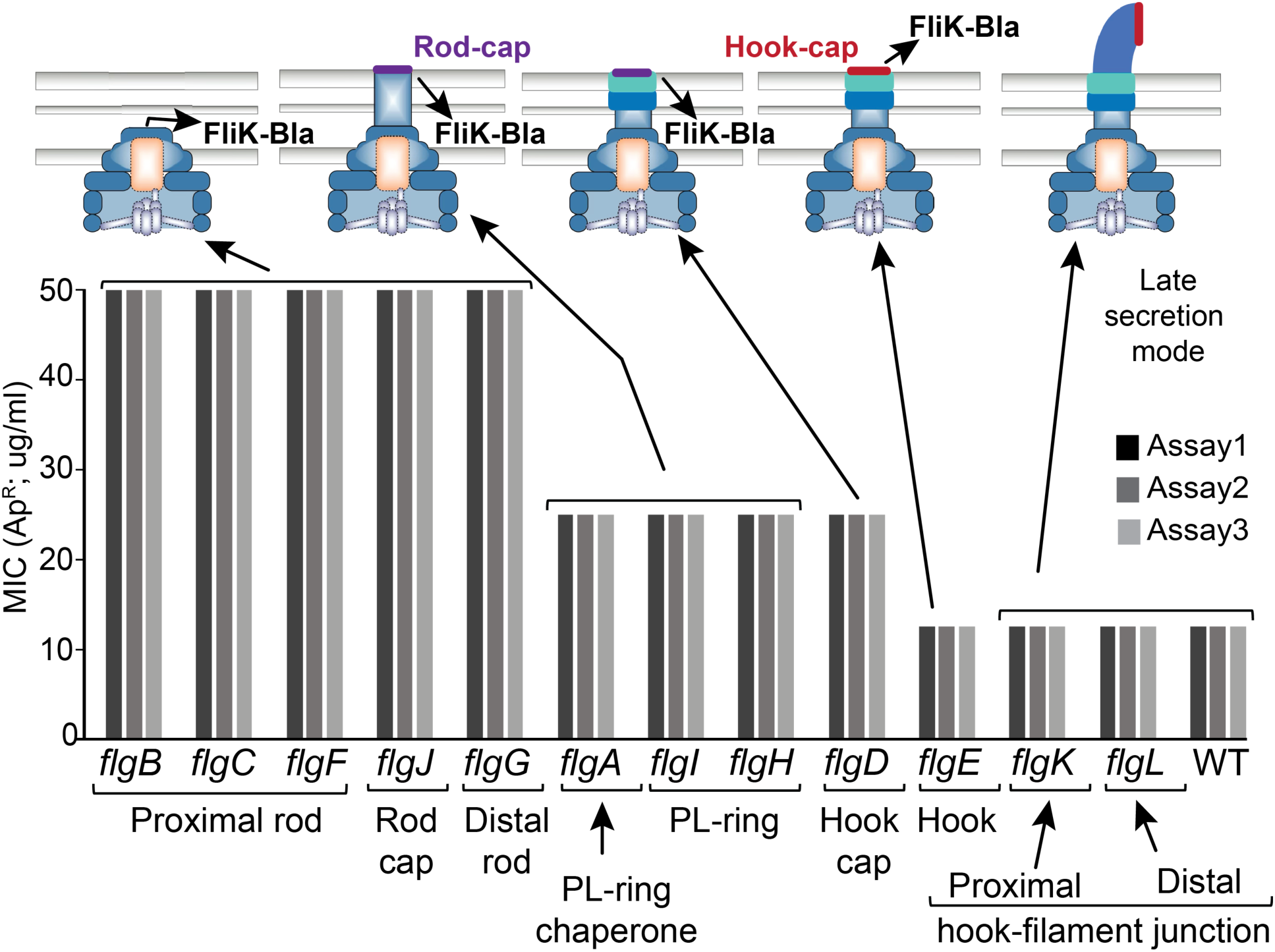
FliK-Bla secretion in different rod-hook mutant backgrounds. Levels of FliK-Bla secretion into the periplasm were assayed in mutants defective in each step of HBB assembly. The MIC to Ap was determined in triplicate for each mutant background. MIC levels were highest in mutants that are unable to polymerize a structure that reaches the outer membrane (*flgB*, *flgC*, *flgF*, *flgJ*). Mutants defective in PL-ring assembly (*flgA*, *flgH*, *flgI*) and the hook cap structural gene (*flgD*) resulted in reduced MIC levels suggesting an inhibition of FliK-Bla secretion at this stage of HBB assembly. Once outer membrane penetration is achieved a further MIC reduction is observed, which likely represents the amount of FliK-Bla that enters the periplasm prior to outer membrane penetration (note that FliK-Bla secreted outside the cell does not fold properly and does not give ApR).

### Removal of the FlhB_CCD_ is dependent on FliK and coupled to HBB completion

As discussed above, our results suggested that FlhB_CCD_ may be physically removed from the HBB during the specificity switch flipping process. To test this idea and whether FliK is required to remove the FlhB_CCD_ upon HBB completion, we developed a system to synchronize expression of the flagellar regulon and thus flagellar assembly. The *Salmonella* regulatory hierarchy is activated by a master regulatory complex FlhD_4_C_2_ that directs RNA polymerase to transcribe the HBB (Class 2) genes. The chromosomal *flhDC* genes were placed under control of a tetracycline (Tc)-inducible promoter (28), and using chromosomally inserted P*_fliF_*-*yfp* and P*_fliC_*-*cfp* indicators (above) for Class 2 and Class 3 flagellar gene expression, respectively, we observed that in an exponentially growing culture Class 2 transcription initiated ∼15 minutes after addition of Tc, and Class 3 transcription began at ∼40 minutes (**Figure 5A**). Using antibodies directed to the FlhB_CCD_, we observed the appearance of cleaved cellular FlhB_CCD_ by 20 minutes and disappearance by about 40 minutes following *flhDC* induction (**Figure 5B**). FlhB_CCD_ was not detected in the spent growth medium suggesting that it is released into the cytoplasm and degraded (**Supplemental Figure 3**).

**Figure 5.**
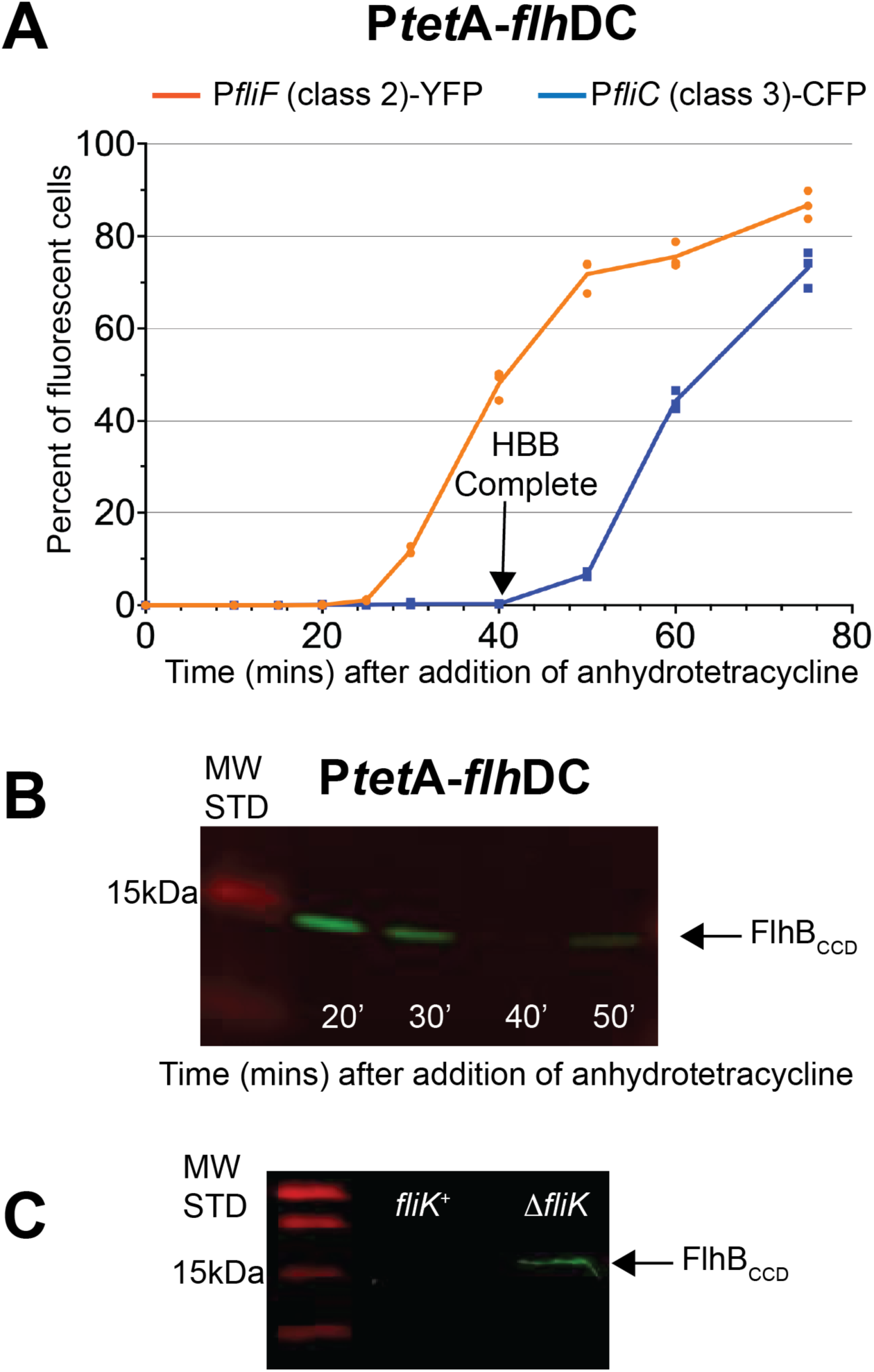
FliK-dependent release of the cleaved C-terminal domain of FlhB. **A.** Percentage of individual cells expressing class 2 (P*_fliF_*-YFP) and class 3 promoter genes (P*_fliC_*-CFP) over-time after induction of the FlhDC master regulator in strain TH27337 (*attB*::Cm^R^-P*_fliF_*-YFP *argW*::zeo^R^-mCherry Δ*galK*::Ap^R^-P*_fliC_*-CFP P*_flhDC_5451*::TPOP *fljB^enx^ vh2*). **B.** Detection of the cleaved C-terminal domain of FlhB overtime after induction of the FlhDC master regulator in strain TH27122 (Δ*araBAD::rflP^+^* Δ*rflM* TPOP-*flhDC fljB^enx^ vh2*). **C.** Effect of FliK on the detection of the cleaved C-terminal domain of FlhB. A strain expressing FliK (TH27622= P*_flhDC_5451*::TPOP Δ*rflM*8403 *fljB^enx^ vh2*) and a strain deleted for FliK (TH27623 =P*_flhDC_5451*::TPOP Δ*rflM8403* Δ*fliK*6140 *fljB^enx^ vh2*) were grown to OD 0.5, at which point anhydrotetracycline was added to the cultures. After 30 min in the presence of anhydrotetracycline, the cells were washed, resuspended in lysis buffer and grown for 30 min before detection.

To determine whether removal of FlhB_CCD_ was dependent on FliK, the same experiment was performed with and without FliK. In addition, at 30 minutes following *flhDC* induction, the cells were washed to remove Tc inducer and thus prevent continued expression of the flagellar regulon. The cells were allowed to grow an additional 30 minutes and analyzed for presence of the FlhB_CCD_. In a *fliK*^+^ background, the FlhB_CCD_ was not detected due to its presumed degradation, but it was present in an isogenic strain deleted for *fliK* (**Figure 5C**). Thus, FlhB autocleavage is not dependent on FliK, but degradation of FlhB_CCD_ is FliK-dependent. These findings support the notion that after HBB completion FliK is required for FlhB_CCD_ degradation. It remains unclear whether FlhB_CCD_ degradation occurs while still bound or after release from the HBB.

### Amino acid substitutions predicted to destabilize FlhB_CCD_ allow FliK-independent switching to late secretion

FliK null mutants are non-motile since they do not proceed to late secretion mode and so have no flagellar filament, and in previous studies motile revertants of a *fliK* null mutant strain (FliK-bypass mutants) resulted in the formation of external polyhook-filament structures (22; 35; 57). DNA sequence analysis revealed these mutants to have FlhB_CCD_ alterations in the AA 274 to 323 region as follows: substitutions S274F, G293V and R, A298V and T, R302P, V310F and Y323D; frameshift and stop codon alleles W348fs, K351-TAG, W353-TAG, and P362fs; a two-residue deletion Δ(G358-Q359; and a change of the TAA stop codon to Y386 which adds 13 amino acids to the C-terminus. The FlhB_CCD_ encompasses 116 residues extending from residue P270 to G385, and a hydrophobic pocket has been identified in it as the docking site for early secretion substrates (6; 47). This pocket contains the side chains of residues A286, P287, A341 and L344. None of the above FliK-bypass amino acid alterations are in the FlhB_CCD_ pocket, and they all secrete both early and late substrates. We propose that these FliK-bypass mutant changes destabilize the folded structure of FlhB_CCD_, so that it fails to function and/or is degraded (above). Autocleavage is not impaired in these mutants suggesting they assemble correctly and FlhB_CCD_ release is necessary before degradation.

Since Fluke also affects switch flipping, we suspected that additional bypass alleles might be obtained in the absence of both FliK and Fluke. Colonies from an overnight streak of strain TH29848 *flhD*C\** Δ*fliK9249* Δ*flk-7755* on L plates were picked and used to inoculate soft-agar motility plates. After one to two days of incubation, motile mutants arose as flairs of swimming cells extending from the site of inoculation (**Fig 6A)**. Twenty independent inoculations all gave rise to substitutions in the FlhB_CCD_ domain that changed seventeen different amino acids. These included 13 new alleles (V260G, S274P, V275L, V289E, ΔG295, R300P, A301V, G305S, V310G, Q349-TAG(Stop), Δ(K351-L355), R354L and Q359-TAG(Stop)) and four previously identified alleles (S274F, G293R, A298V, W353-TAG, above). Upon prolonged incubation several motile flairs gave rise to secondary mutations resulting in enhanced motile phenotypes. All the secondary substitutions with enhanced motile phenotypes were also in the FlhB_CCD_ domain with the following multiply substituted alleles: S274F+V257L, S274F+Δ(L350-R354), S274F+V310G, S274F+G294V+E314K, G305S+P370T, G305S+A319V, G305S+A294P, W353-TAG(Stop)+K292R, and R354L+A339V. The FliK/Fluke-bypass alleles were mapped onto the FlhB_CCD_ structure (**Figure 6B and 6C**), and all changes are predicted, based on the amino acid changes, to result in a destabilization of the FlhB_CCD_ folded structure. This further supports the notion that FlhB_CCD_ inactivation by removal, unfolding and/or degradation is important in flipping secretion switch to specificity.

**Figure 6.**
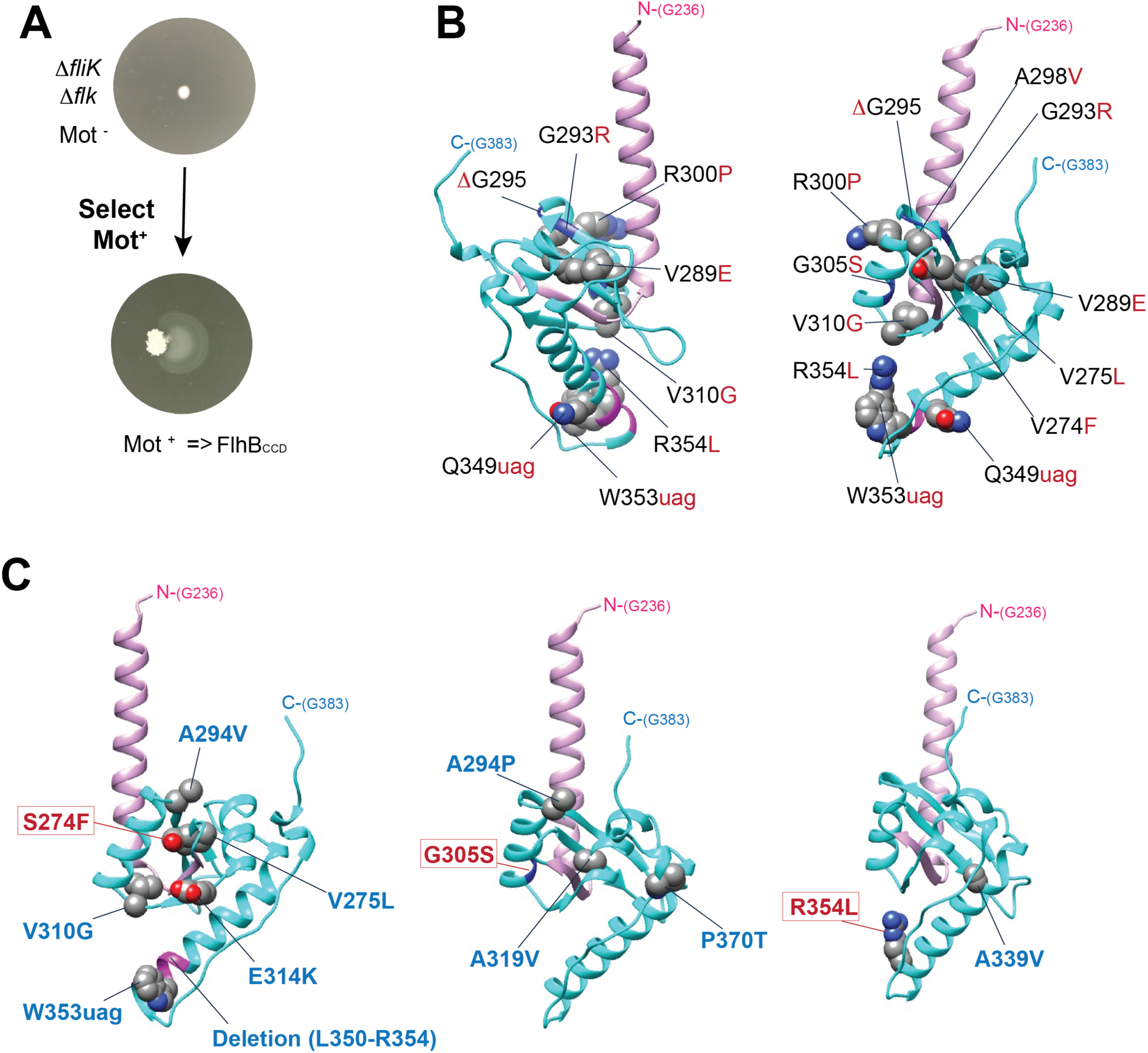
Selection for motile revertants of Δ*fliK* in a Δ*flk* mutant background result in mutants in the C-terminal domain of FlhB. **A.** Motile suppressors selection. **B.** Single mutations obtained in FlhB_CCD_ (backbone colored in cyan). The suppressor mutations are shown in red. The lavender helix includes residue G331 through N269 FlhB_N_ that are located just before the site of autocleavage site between N269 and P270. **C.** Secondary suppressor mutations with increased motility phenotypes isolated from initial motile suppressors (S274F, G305S and R354L, colored in red) isolated in the Δ*fliK* Δ*flk* double mutant background are shown in blue.

### Alterations in the secretion pore allow FliK-dependent switching in the absence of FliK-tape measure length measurement

The above selections for mutants that allow late secretion in the absence of FliK and Fluke found only changes that destabilize or remove FlhB_CCD_, but we reasoned that other rarer changes and thus other involved proteins might be found if these could be avoided. These *flhB* mutations are expected to be recessive to the wild-type *flhB*^+^ allele (and indeed deletion of FlhB_CCD_ is recessive), so we repeated this selection with two copies of the *flhB* gene present (one is expressed from the *araBAD* promoter (P*_araBAD_*-*flhB*^+^); strain TH27965, **Figure 7A**). Both copies of *flhB* would have to be altered to destabilize all FlhB proteins, and the frequency of such double mutants should be very low. The frequency of mutations that allow successful FlgM-Bla secretion (Ap^R^) in this strain was indeed 100-fold lower than in the selection strain carrying a single copy of the *flhB* gene, but the rare mutants obtained in this selection yielded only seven different mutations from 40 independent selection experiments. One substitution in the FliK_C_ switch domain, P296L was not unexpected since FliK is a known switch participant and P296S and P296T substitutions in FliK have been previously identified as intragenic motile suppressors of a non-motile *fliK* mutant that lacks amino acids 271-400 (31). The other six mutations were surprising in that they altered proteins that make up the secretion pore: FliP(I195N), FliQ(G32D), FliR(Q210-TAG), FliR(S212-TAG), FliR(V215D) and FliR (duplication of codon T221). Secretion of FlgM-Bla in the strains carrying these mutant secretion pore proteins is dependent on the absence of Fluke, so Fluke can inhibit the switch to late secretion in the pore protein mutants (**Figure 7C**).

**Figure 7.**
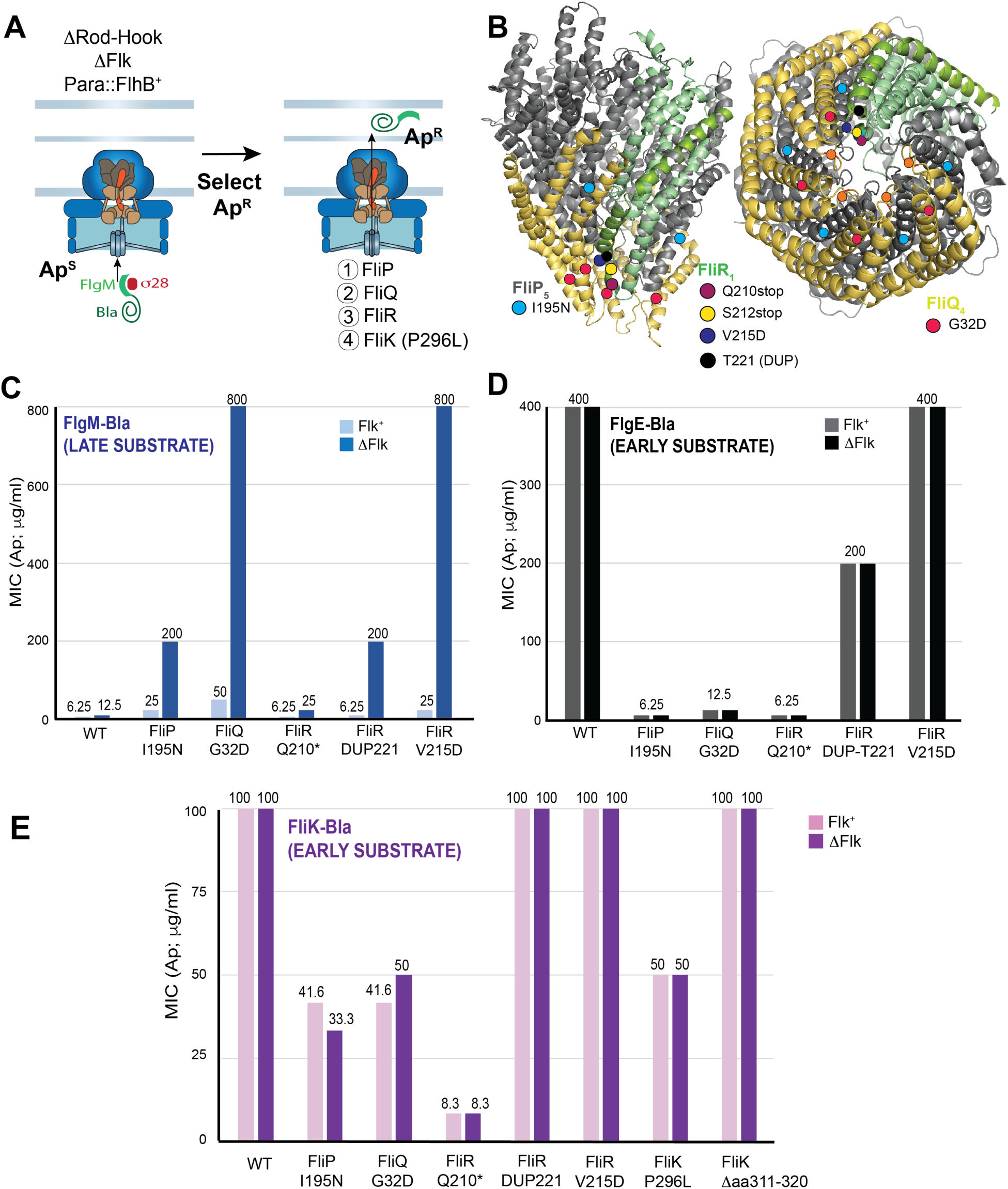
Late-secretion FlhB-bypass mutants obtained in the secretion pore (FliPQR) and FliK. **A.** Selection strain is missing the early rod-hook secretion substrates and Fluke with a second copy of *flhB*^+^ under control of an arabinose promoter. Selection for FlgM-Bla secretion with both *flhB*^+^ genes expressed yields rare PPBS-Ara-Ap15^R^ FlhB-bypass alleles in the core secretion pore genes *fliP*, *fliQ* and *fliR* and the *fliK* ruler gene. **B.** FlhB-bypass mutations in *fliP*, *fliQ* and *fliR* are all located to the bottom of the pore through which secreted substrates pass; veiw on right is from the cytoplasmic side of the structure. **C.** Secretion of FlgM-Bla in the FlhB-bypass mutant backgrounds exhibit is inhibited in the presence of Fluke. **D. & E.** The effect of FlhB-bypass mutations on early substrate FlgE-Bla (**D.**) and FliK-Bla (**E.**) secretion is essentially Fluke-independent. Strains used in this assay are listed in Table S4.

To test the functionality of the mutant pores, we measured the effect of these mutations on early substrate secretion using FlgE(Hook)-Bla (**Figures 7D**) and FliK-Bla (**Figures 7E**) early secretion reporters in strains defective in rod-hook assembly. Most of these mutations caused significant reduction in FlgE-Bla and FliK-Bla secretion indicating that the mutant pores are substantially defective in early secretion (we suspect that the FliR(V215D) allele, which did not have an effect in these measurements, may also be partially secretion defective, but not enough to detect in our assay). These measurements were not affected by the presence or absence of Fluke as is expected for early secretion. These experiments allow us to propose a novel mechanistic model where a reduction in the rate of FliK secretion that is coupled to length measurement allows FliK_C_ to have a productive interaction with FlhB_CCD_ and thus cause T3S specificity switch flipping.

We were puzzled that alleles defective in the secretion pore proteins FliP, FliQ and FliR result in the switch to late substrate (FlgM-Bla) secretion until we discovered that introduction of a *fliK* deletion prevented FlgM-Bla secretion in these strains (**Supplemental Figure 5**). That is to say that in the absence of a rod-hook structure, whose length is normally measured by FliK resulting the switch at a rod-hook length of ∼80 nm, FliK is flipping the secretion-specificity switch in the FlhB-bypass mutant strains without any structure to measure! Thus, FliK is still required to catalyze the removal of FlhB_CCD_ and Fluke as well as the transition of FlhA from the closed to open form. Since the speed of FliK secretion determines whether FliK_C_ can flip the secretion-specificity switch (18), we reasoned that the mutations in FliP, FliQ and FliR result in slow FliK secretion that allows FliK_C_ to flip the switch.

We also suspected that the ability of the FliK P296S and P296T substitution to suppress the motility defect of the *fliK* mutant lacking amino acids 271-400 (31) might be due to a reduced rate of FliK secretion and the same could be true for the P296L substitution isolated above. If FliK(P296L) is secreted at a slower speed it could result in a premature T3S switch earlier in HBB assembly that wild-type FliK. This was tested by placing the *flhDC* master operon under a Tc-inducible promoter and determining if the T3S switch occurs earlier than with FliK^+^. The T3S switch to late mode results in FlgM secretion, which was assayed using a *lux* operon promoter expressed from the Class 3 *fliC* promoter. As shown in **Figure 8**, the strain expressing the FliK(P296L) allele resulted in flipping of the T3S switch substantially earlier than with FliK^+^, which is consistent with the P296L substitution resulting in a reduced rate of FliK secretion. As a control, we showed that a mutant FliK deleted for amino acids 311-320, that was previously shown to be secreted but unable to catalyze the T3S switch did not turn on the *lux* genes in this strain (38). We propose that the P296L substitution affects the ability of FliK to pause during secretion and interact to flip the switch (see Discussion).

**Figure 8.**
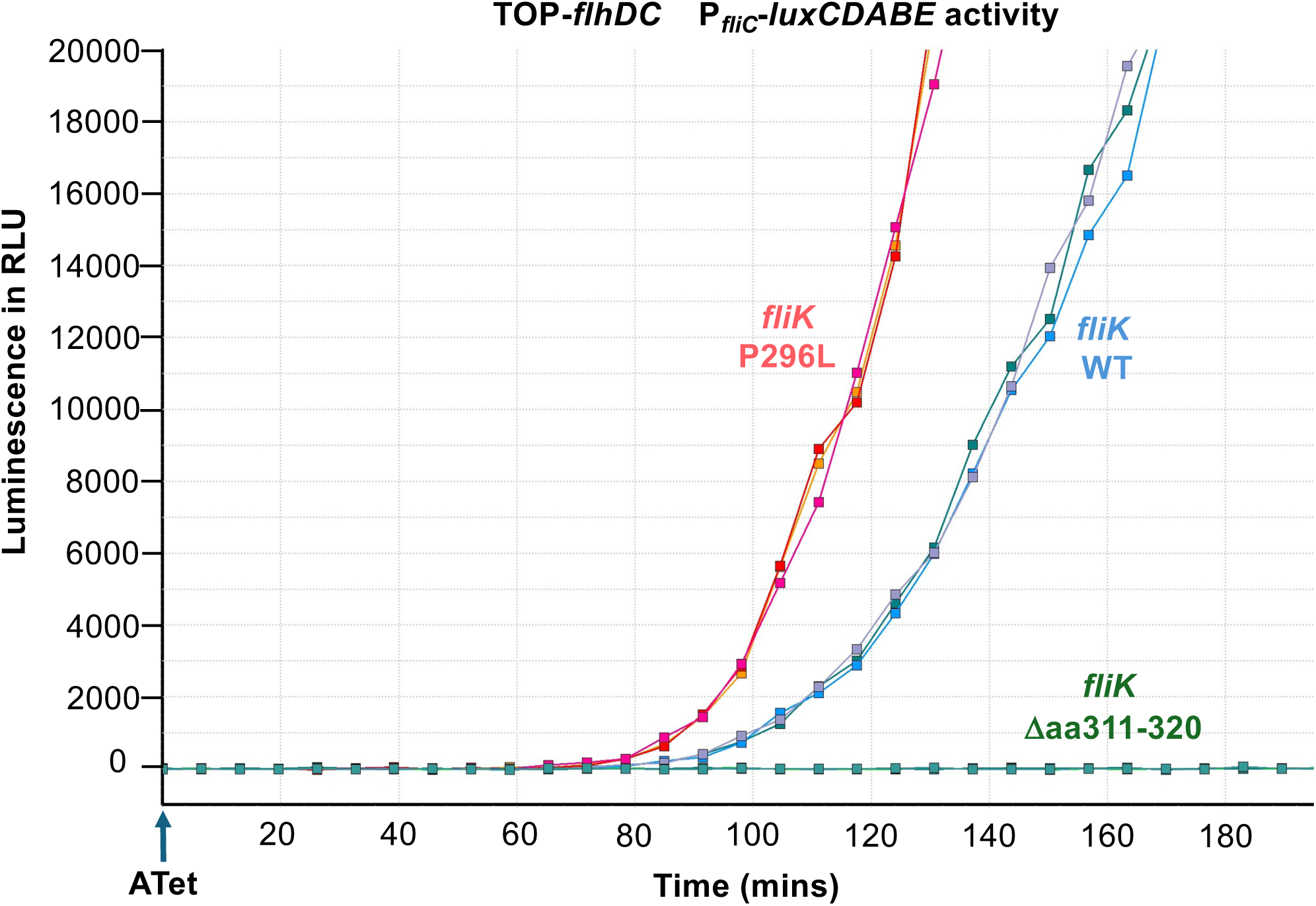
Effect of *flik* alleles on the timing of T3S Specificity Switching. Class 3 promoter activity was monitored using a *fliC*–*luxCDBAE* luciferase reporter in strains expressing wild-type *fliK* (TH30413), *fliK* Δaa311–320 (TH30414), or *fliK* P296L (TH301415) under a tetracycline-inducible *flhDC* promoter. Cultures were induced at OD₆₀₀ = 0.5 with anhydrotetracycline and luminescence was measured every 5 min in a plate reader. Data represent the mean of four technical replicates from three independent biological replicates; colors indicate individual biological replicates for each strain.

## Discussion

Bacterial flagella are large macromolecular assemblies whose construction requires precise coordination between secretion, assembly and gene regulation. A defining feature of flagellar biogenesis is the irreversible switch in type III secretion (T3S) specificity that terminates hook growth and initiates filament assembly. Although the molecular ruler FliK and the core secretion component FlhB have long been known to control this transition, how hook-length sensing is converted into a stable change in secretion specificity has remained unresolved. The results presented here identify FlhB_CCD_ and Fluke as inhibitory components that actively maintain the flagellar T3S apparatus in early secretion mode and demonstrate that their FliK-dependent removal irreversibly triggers the specificity switch.

Classical genetic studies identified mutants lacking FliK as defective in secretion switching, which produced elongated polyhooks (34; 45; 52). These observations established FliK as the catalyst of the switch but did not explain how switching is prevented prior to hook completion. Our findings support a model in which early secretion is not simply the default state of the apparatus but is actively enforced by inhibitory components. FlhB_CCD_ and Fluke function as molecular locks that must be removed to permit the transition to late secretion and filament assembly.

FliK contains two functionally: an N-terminal ruler domain that determines hook length (39; 51) and a C-terminal specificity switch domain that interacts with FlhB (31; 38). Alterations in the length of the ruler domain proportionally alter hook length (51), supporting a physical ruler mechanism. Our results further refine this model by showing that the rate of FliK secretion is a critical determinant of productive switching. When FliK is secreted rapidly, as occurs in the absence of a rod or hook structure (18), interaction between FliK_C_ and FlhB_CCD_ is inefficient and switching does not occur. In contrast, conditions that slow FliK transit through the secretion channel, either during normal hook elongation or in secretion rate bypass mutants, permit productive interaction and triggers switching even in the absence of hook assembly.

Structural modeling and mutational analysis support a mechanism in which the terminal residues of FliK destabilize FlhB_CCD_, promoting its unfolding and removal from the secretion apparatus. Our data indicate that a folded region within FliK_C_ contributes to the timing of this interaction, likely by introducing a transient secretion pause that allows the C-terminal residues to engage FlhB_CCD_ (**Figure 9A**). This model is consistent with the observation that FliK variants lacking this folded region are secreted efficiently but fail to induce switching, uncoupling secretion from specificity control (38).

**Figure 9.**
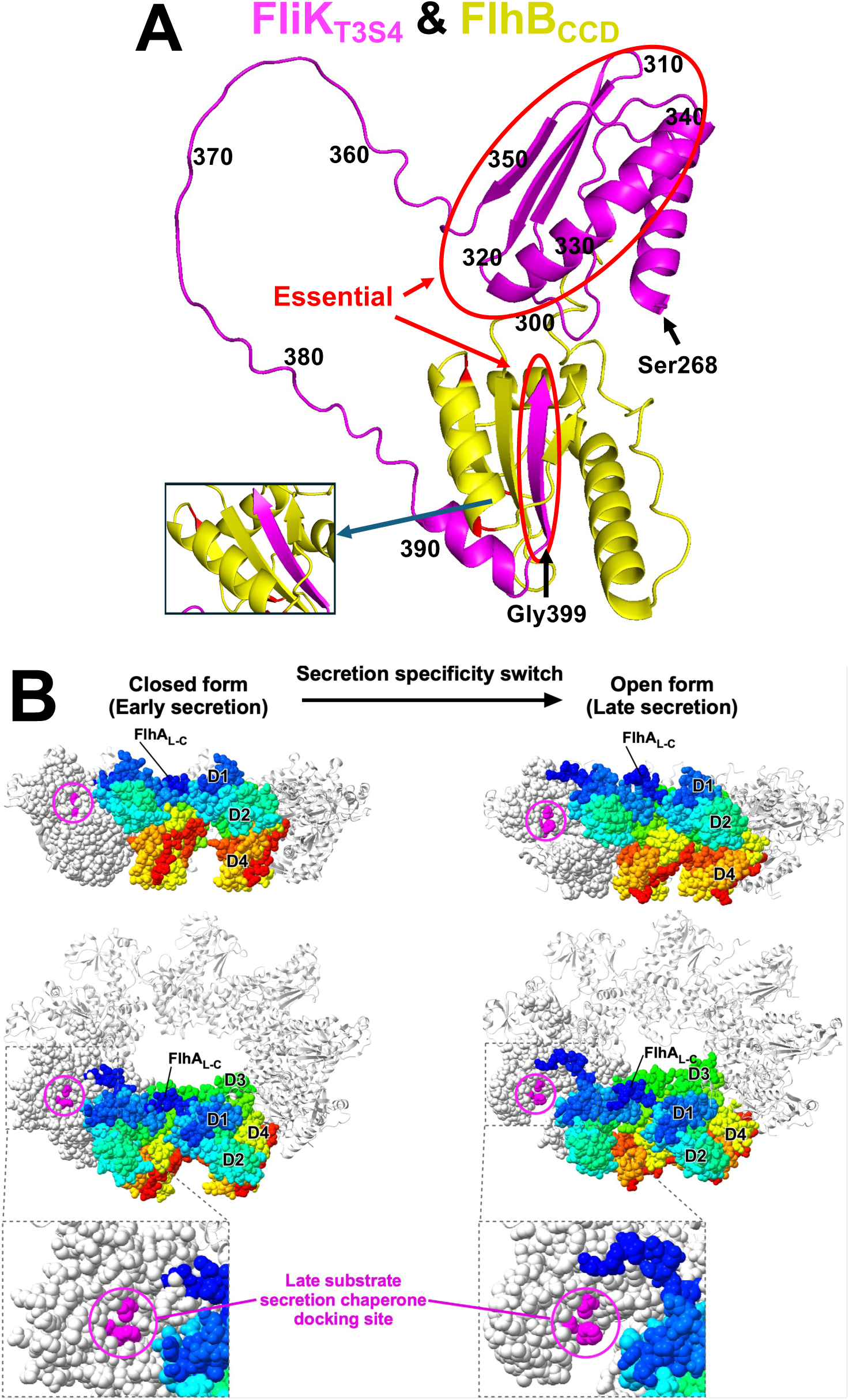
A. Alpha-Fold model of the interaction between FlhB_CCD_ and FliK C-terminal T3S4 domain. Structural models of the C-terminal domains of FliK (residues 269–end; colored in magenta) and the cleaved form of FlhB (residues 270–end; colored in yellow) were generated using the AlphaFold3 prediction server (1). **B. Structural models of the open and closed forms of the FlhA_C_ ring.** The closed and open ring models were generated by fitting domain D1 of the open (PDB ID: 3A5I) and closed (PDB ID: 3MYD) conformations onto the corresponding domain of each FlhA_C_ subunit in the cryo-EM structure of the FlhA_C_ ring (PDB ID: 7AMY). Two FlhA_C_ subunits are color-coded from blue to red, following the rainbow spectrum from the N-terminus to the C-terminus, while the remaining subunits are shown in light grey. One FlhA_C_ subunit is displayed in CPK representation. Residues involved in the interaction with flagellar export chaperone in complex with their cognate filament-type substrates are highlighted in magenta.

In addition to FlhB_CCD_, our data implicate Fluke as a second inhibitory component that prevents premature switching. Genetic assays suggest that Fluke functions at the level of FlhA, a central component a secretion apparatus that undergoes a conformational change during switching. Previous work has shown that FlhA adopts a closed conformation prior to switching, occluding the docking site for late substrate-chaperone complexes, and transitions to an open conformation following switching (**Figure 9B**) (24). Our results suggest that Fluke stabilizes the closed conformation of FlhA, thereby reinforcing the early secretion state. Removal of Fluke increases the probability that FliK-mediated switching occurs, particularly under conditions in which secretion dynamics are altered at the rod to hook transition in flagellar assembly.

Our findings support a model in which secretion specificity switching is governed by the coordinated removal of inhibitory components in response to assembly state completion. As illustrated in **Figure 10**, completion of hook assembly positions a secreting FliK molecule such that its C-terminal domain can productively interact with FlhB_CCD_. A secretion pause, facilitated by unfolding within FliK_C_, allows the terminal residues of FliK to destabilize and eject FlhB_CCD_ from the apparatus. This event, coupled with destabilization of Fluke-dependent inhibition at FlhA, drives the transition from early to late secretion by exposing the secretion-chaperone docking site required for filament assembly.

**Figure 10.**
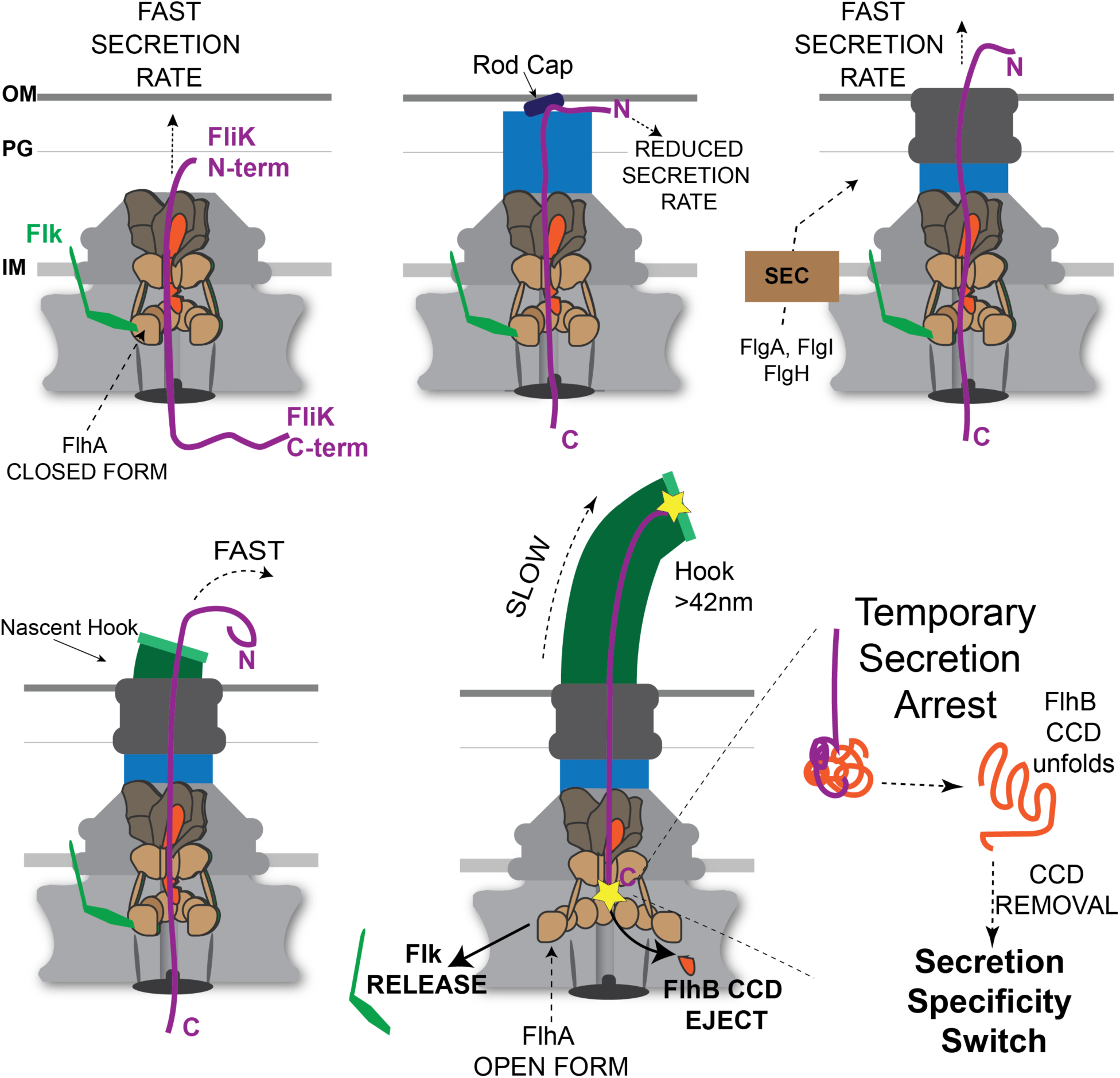
FliK temporary secretion arrest model. FliK is slow to reach the tip of the growing flagellar organelle, but once the N-terminus of FliK exits the structure it is secreted at a relatively fast rate (18). Rod terminates at the outer membrane. PL-ring formation and outer membrane (OM) penetration are an uncoupled process to rod completion. Figure 4 data suggest FliK secretion into the periplasmic space is impaired with the rod cap against the OM. This appears to be a critical step in HBB assembly, and the system has evolved to utilize two proteins to protect the structure at this stage of assembly. A periplasmic protein FlhE helps to maintain the rod cap in place. Cells lacking FlhE have a high rate of rod cap loss replaced by the hook cap prior to PL-ring assembly resulting in periplasmic flagella (21). We propose that the Fluke protein stabilizes FlhA in its closed conformation such that a reduced rate of FliK secretion upon rod completion does not prematurely flip the secretion specificity switch. Following PL-ring formation and OM penetration, FliK is again secreted at a fast rate once the N-terminus of FliK exits the structure. However, the slow transit to reach the tip is essential but not solely responsible to flip the switch. The C-terminal FliK T3S4 domain contains a protein fold encompassing residues 301-350 that is also essential to flip the switch. We propose that this fold is required to facilitate a temporary secretion arrest and provide a final time element essential for the last 5 amino acids of FliK to interact with FlhB_CCD_ and flip the switch.

This dual inhibitor mechanism ensures that secretion switching occurs only after the hook reaches its optimal length, preventing premature filament assembly and coordinating secretion with transcriptional activation of late flagellar genes. More broadly, our work reveals that that the flagellar T3S system employs active maintenance of early secretion through inhibitory components, rather than relying solely on passive sensing of assembly completion. This principle may extend to other regulated secretion systems in which irreversible transitions must be tightly coupled to structural assembly checkpoints.

## Materials and Methods

### Bacterial strains and growth conditions

Bacterial strains used in this study are derived from *Salmonella enterica* serovar Typhimurium wild-type strain LT2 and are listed in Table S1. Bacteria were grown in Lysis Broth (LB: 10 g/l Tryptone, 5 g/l Yeast extract, 5 g/l NaCl) in liquid culture with shaking or on solid media (LB supplemented with 12 g/l Apex agar). Antibiotics were added to LB media for selections or plasmid retention at the following final concentrations: 100 µg/ml sodium ampicillin (Ap), 12.5 µg/ml chloramphenicol (Cm), 15 µg/ml tetracycline-HCl (Tc), 50µg/ml kanamycin sulfate (Km), 100 µg/ml carbenicillin. Gene expression from the TPOP or arabinose promoter was induced with 125 ng/ml anhydrotetracycline (ATc) or 0.2% arabinose, respectively. Selection for tetracycline-sensitive mutants was done on fusaric acid plates (12 g/l Apex agar, 5 g/l Bacto tryptone, 5 g/l yeast extract, 10 g/l NaCl, 10 g/l NaH₂PO₄·H₂O, 12 µg/ml fusaric acid, 0.5 µg/ml ATc). Selection for Ap^R^ mutants was done on PPBS medium (17 g/l Bacto peptone, 3 g/l Bacto-proteose peptone, 1.5 g/l Difco bile salts #3, 10 g/l NaCl, 11 g/l Apex agar) containing 15 µg/ml sodium ampicillin (PPBS-Ap15).

### Strain construction

Strains used in this study were engineered by generalized phage transduction and/or chromosomal recombineering using the λ-Red recombineering system. The generalized transducing phage of *Salmonella enterica* serovar Typhimurium, P22 *HT105/1 int-201*, was used to transfer genetic alleles between strains (14). For P22 transductions, 100 µl of 10⁶ plaque-forming units (PFU)/ml of P22 transducing lysate grown on donor strains was mixed with 100 µl of an overnight culture of recipient cells (2 × 10⁸ cells/ml). When necessary, mixtures were incubated at 37°C or 30°C for 30 min to allow phenotypic expression before plating on selective media.

Selection for replacement of Tn*10dTc* alleles or chromosomal *tetRA* cassette insertions by P22 transduction was performed as follows. Fresh P22 lysate grown on the donor strain was diluted to ∼10¹⁰ PFU/ml in sterile saline (0.85% NaCl). 0.1 ml of the diluted phage was mixed with 0.1 ml of a freshly saturated recipient culture (1–5 × 10⁹ cells/ml) and incubated at 37°C without shaking. After 30 min, 20 µl of the phage–cell mixture was diluted into 5 ml sterile saline, and 0.2 ml of the diluted suspension was plated onto fusaric acid plates. A control containing 0.1 ml saline and recipient cells only was included to determine the frequency of spontaneous Tcˢ mutants. Typically, we observed ∼100-fold more Tcˢ colonies on transduction plates than on control plates. Tcˢ colonies were purified on green indicator plates (14) and genetic alleles were verified by PCR. Chromosomal gene editing (recombineering) was performed using the helper plasmid pSim5 (Cmᴿ) to insert and remove a *tetRA* cassette derived from transposon *Tn10*, following an established protocol (26). Replacement of the *tetRA* cassette by λ-Red recombination was achieved by selecting for fusaric acid resistant mutants, as described above. Background spontaneous Tcˢ mutants conferring fusaric acid resistance were minimized by diluting electroporated cells 10-100-fold prior to selection at 42°C. Genetic alleles were verified by DNA sequencing. Phusion polymerase was used for all λ-Red PCR-generated constructs, whereas Taq polymerase was used for PCR verification of construct size and sequencing reactions.

### Construction of *flhB*_Δ__CCD_

An flhB allele lacking the cleavable C-terminal domain of FlhB was generated as follows. A *tetRA* cassette was amplified from genomic DNA of strain TH408 using primers 1874 and 1910 and inserted into strain TH27083 by λ-Red recombination, selecting tetracycline resistance (Tc^R^) to yield strain TH23957. The *tetRA element* was then replaced with a clean deletion using a fill-in reaction between primers 1911 and 1912 and λ-Red recombination, selecting for Tc^S^ on fusaric acid plates. This yielded to strain TH23959. The resulting *flhB*_ΔCCD_ deletion (Δ amino acids P270-G283) preserved the ribosome-binding site of *flhA* and the last 9 codon sequence of *flhB*. Complementation was achieved using a plasmid expressing *flhB*. The *flhB* allele was introduced into strains of interest by P22 phage transduction.

### Selection for late (FlgM-Bla) secretion in the absence of Fluke and hook-rod proteins yielding mutations in the FlhB_CCD_

β-lactamase (lacking its Sec-dependent signal sequence) fused to the C-terminus of FlgM (FlgM-Bla) (9) was used to select for late substrate secretion into the periplasm. Strain TH25226 (*flgM6427*::*bla* Δ*flgB-L8735 flhD8070 flhC8092 fliA5225* Δ*fliB-T7771* Δ*flk-7755 fljB^enx^ vh2*) lacks all rod-hook structural genes except fliE (*flgB-L8735=*Δ*flgBCDEFGHIJKL)*, all late secretion substrates except FlgM-Bla (Δ*fliB-T7771=* Δ*fliBCDST, fljB^enx^ vh2=* Δ*fljAB-hin)*. Strain TH25226 is also deleted for the *flk* gene and contains the protease resistant alleles for the proteins of the flagellar master regulatory complex FlhD_4_C_2_ that increases the number of hook basal bodies/bacteria from ∼4 to ∼12 (termed the *flhD*\**C*\* background) (16). From each of ten independent overnight cultures of strain TH25226, 100 μl was plated on PPBS-Ap15 plates and incubated at 37°C. The following day a thin film of growth appeared on the plates. The were replica printed a second time on PPBS-Ap15 plates and following overnight incubation at 37°C only single colonies appeared at a frequency of ∼5 x 10^-7^ and the background growth was no longer present. Ap^R^ colonies from each independent selection plate were purified on non-selective LB medium and the Ap^R^ phenotypes were retested on PPBS-Ap15 plates. Using linked transposons to the *flg*, *flh* and *fli* regions, Ap^R^ mutations were mapped. P22 transduction frequencies to linked transposons were used to estimate specific chromosomal location of mutant alleles (49). Targeted DNA regions were amplified by PCR and subject to DNA sequence analysis. To identify possible *flhA* and *flhB* mutants, we also pooled the Ap^R^ colonies form the above selection plates and transduced the pooled cells into strain TH26992 (*flgM6427*::*bla* Δ*flgB-L8735 STM1911*::Tn*10d*Tc Δ*flhBAE7670*::FCF *flhD8070 flhC8092 fliA5225(H14D)* Δ*fliB-T7771* Δ*flk-7755 fljB^enx^ vh2*) selecting for replacement of the *STM1911*::Tn*10d*Tc allele on Tc^S^ selection plates. The Tc^S^ transductants were screened for replacement of the Δ*flhBAE7670*::FCF allele by their Cm^S^ phenotype. The alleles linked to the *flhBAE* region (Tc^S^ Cm^S^ transductants) that inherited the Ap^R^ phenotype were subject to DNA sequence analysis of the *flhBA* loci.

### Selection for late (FlgM-Bla) secretion in the absence of the cleavable C-terminal domain of FlhB (FlhB_CCD_) yielding mutations in the *flk* and *secY* loci

Strain TH24722 (*flgM6427::bla* Δ*flgB-L8735 flhB8612*(ΔCCD) *flhD8070 flhC8092 fliA5225*(H14D) Δ*fliB-T7771 fljB^enx^ vh2*) was constructed so that it was (i) deleted for the *flhB* cleaved C-terminal domain *flhB8612*, (ii) deleted for the genes encoding the rod and hook structure and assembly proteins (Δ*flgB-L8735*), (iii) carried the *flhD*\**C*\* alleles (*flhD8070 flhC8092*) and a FliA allele (*fliA5225*) resulting in two-fold increased σ^28^ levels (7), (iv) deleted for late secretion substrates (Δ*fliB-T7771 fljB^enx^ vh2*) and carrying the full length FlgM with a C-terminal fusion of β-lactamase deleted for its N-terminal Sec secretion signal (*flgM6427::bla*). From each 10 independent cultures, ∼5 x 10^8^ cells were plated onto separate PPBS-Ap15 plates. After overnight incubation at 37°C revertants would appear with a thin film of background growth on the plates. The plates were replica printed back onto PPBS-Ap15, which eliminated the background film of cell growth. Ap^R^ mutant colonies appeared at a frequency of ∼10^-6^. All ten original cultures yielded mutants that were Ap^R^ at 30°C and 37°C, but included a mixture of cells that were either Ap^S^ or Ap^R^ at 42°C. One 30°C-37°C-42°C Ap^R^ and one 30°C-37°C Ap^R^ 42°C Ap^S^ colony from each independent plating was kept for linkage analysis. Nine 30°C-37°C-42°C Ap^R^ colonies were linked to the *flk* locus and one to the *secY* locus. Additionally, three independent cultures yielded 37°C-42°C Ap^R^ 30°C Ap^S^ mutants. Two were linked to *SecY* and one to *flk*.

### Selection for late (FlgM-Bla) secretion using a strain lacking Fluke and duplicated for the *flhB* gene yielding mutations in *fliK*, *fliP*, *fliQ* and *fliR*

FlhB-bypass mutants were isolated by repeating the above described Ap^R^ selections in a strain expressing two copies of the *flhB*^+^ gene: one from the normal *flhB* locus and the second from the *araBAD* locis. Strain TH27965 (Δ*araBAD2101*::*flhB*^+^ *flgM6427*::*bla* Δ*flgB-L8735 flhD8070 flhC8092 fliA5225*(H14D) Δ*fliB-T7771* Δ*flk-7755 fljB^enx^ vh2*) was grown to saturation in LB containing 0.2% arabinose and 10^9^ cells from ten independent cultures were plated onto PPBS-Ap15-Ara(0.2%) plates, incubated overnight at 37°C, replicate printed again onto PPBS-Ap15-Ara(0.2%), incubated overnight at 37°C. Ara-Ap^R^ colonies appeared at a frequency of ∼10^-9^. One large and one small Ara-ApR colony was picked from each independent culture and checked for linkage to the *secY*, *flhBAE* and *fliE-fliR* regions. Nine colonies were linked to the *secY* locus and DNA sequence analysis revealed them to include one of the following amino acid substitutions in SecY: G69D(3), A71V(3), G194D(3) and P276S(1). Similar mutants in *secY* of *E. coli* have been reported that allow for secretion of LamB and PhoA lacking their Sec-dependent secretion signal (15). None were linked to the *flhBAE* region, and ten that were linked to the *fliE-R* region were all G32D substitutions in FliQ. The G32D substitution in FliQ was recessive to *fliQ*^+^ expressed from a salicylate(Sal)-inducible *nahG* promoter from plasmid pEM5 (Km^R^ P*_nahG_*-*fliQ*^+^). Because the G32D substitution in FliQ appeared to result from targeting a “hot-spot” for spontaneous mutation, the selection was repeated using strain TH27965 carrying the pEM5 plasmid. Ten independent cultures were grown to saturation in LB-Ara(0.2%)-Km and 10^9^ cells from each culture were plated onto PPBS-Ap15-Ara(0.2%)-Sal(10µM)-Km plates. Eight plates yielded Sal-Ara-Ap^R^ mutants and those linked to the *fliE-R* region were sequenced and included mutations in *fliK*(P296L), *fliP*(I195N) and *fliR*(Q210-TAG, V215D and duplication of the T221 codon). We also specifically targeted the alpha helix that began with the G32 position in case there was another important genetic target there. Codons 31 through 50 of *fliQ* was replaced with a *tetRA* cassette followed by replacement of the *tetRA* cassette with an oligonucleotide covering the region that had on average one base substitution in the codon 31 through 50 region per oligo as described (26). Tc^S^ cells were pooled and the mutagenized region was moved into the TH27965 strain by P22 transduction and screened for growth on PPBS-Ap15-Ara(0.2%). Two substitutions were identified in FliQ, T48M and G32E resulting in an FlhB-bypass phenotype.

### Selection for FliK by-pass motile revertants in a *flk* null background

Strain TH29848 (*flhD8070 flhC8092* Δ*fliK9249* Δ*flk-7755*) that lacks FliK and Fluke was used to select for motile, FliK-bypass mutants in FlhB. Colonies from an overnight streak of TH29848 on L plates were pick by toothpick and used to inoculate soft-agar motility plates. After one to two days on incubation motile revertants arose as flairs of swimming cells extending from the site of inoculation. From twenty independent inoculations, thirteen different amino acid substitutions in the FlhB_CCD_ domain were obtained: V260G, S274F, S274P, V275L, V289E, G293R, ΔG295, A298V, R300P, A301V, G305S, V310G, Q349-TAG(Stop), W353-TAG(Stop) Δ(K351-L355), R354L, and Q359-TAG(Stop). Upon prolonged incubation several motile flairs would give rise to secondary mutation resulting in more enhanced motile phenotypes. All secondary substitutions with enhanced motile phenotypes were also in the FlhB_CCD_ domain resulting in the following multiply substituted alleles: S274F+V257L, S274F+Δ(L350-R354), S274F+V310G, S274F+G294V+E314K, G305S+P370T, G305S+A319V, G305S+A294P, W353-TAG(Stop)+K292R, and R354L+A339V.

### Fluorescent transcriptional reporters to flagellar class 2 and class 3 promoters

Transcriptional reporters for flagellar class 2 and class 3 promoter expression utilized promoters from the *fliF* (class 2) or *fliC* (class 3) loci. Reporter fusions to either yellow fluorescent protein (mVenus NB) (YFP) or cyan fluorescent protein SCFP3A (CFP) were constructed that included the ribosomal binding site of gene 10 of phage T7 as described (30). The flagellar class 2 and class 3 transcriptional reporter cassettes were first PCR amplified from *E. coli* strain MG1655 MGR FC (30), using primers 8283/8284 for the P*_fliFcoli_*-CFP-Ap^R^ cassette and primers 8285/8286 for the P*_fliCcoli_*-YFP-Km^R^ DNA cassette. These cassettes were introduced respectively into the *galK* and in the *attB* locus of the *Salmonella* chromosome by λ-Red recombineering, selecting Ap^R^ for inheritance of the P*_fliFcoli_*-CFP-Ap^R^ cassette, and Km^R^ for inheritance of the P*_fliCcoli_*-YFP-Km^R^ cassette. The close chromosomal proximity of the two *loci* minimizes the impact of copy number variation during replication. Once integrated into the *Salmonella* genome, *tetRA* elements were inserted in front of the CFP and YFP genes, replacing the E. *coli fliF* and *fliC* promoters. This was done using λ-Red and primers 8315/8331 and 8309/9083, respectively. The *tetRA* elements were then replaced using α-Red mediated gene editing on Tc^S^ selection medium with the *Salmonella fliF* and *fliC* promoters using primers 8459/8460 (resulting in P*_fliF_-YFP*) and 9142/8458 (for P*_fliC_-*CFP). For ease of genetic manipulations, we also exchanged by λ-Red the kanamycin cassette in the P*_fliF_*_(*Salmonella*)_*-*YFP-Km^R^ with a chloramphenicol marker, amplified from plasmid pKD3 (13), using primers 9505/9506. Once inserted in the *Salmonella* chromosome the *attB*::P*_fliF_*_(*Salmonella*)_-YFP-Cm^R^ and the Δ*galK*::P*_fliC_*_(*Salmonella*)_*-*CFP-Ap^R^ were moved to appropriate strain backgrounds by phage P22-mediated generalized transduction.

### Microscopy analysis of the transcriptional reporters to class 2 and class 3 flagellar genes

Strains carrying the *attB*::P*_fliF_*_(*Salmonella*)_-YFP-Cm^R^ and the Δ*galK*::P*_fliC_*_(*Salmonella*)_*-*CFP-Ap^R^ transcriptional reporters were diluted 1:100 from overnight cultures and grown in LB broth to an optical density (OD₆₀₀) of approximately 1. For the time-course experiment (Figure 5), strain TH27337 (*attB*::Cm^R^-P*_fliF_*-YFP *argW*::zeo^R^-*P_const_*-mCherry Δ*gal<u>K</u>*::Ap^R^-P*_fliC_*-CFP P*_flhDC_5451*::TPOP *fljBenx vh2*) was grown in 25 ml LB from an overnight culture. When the culture reached an OD₆₀₀ of 0.5, expression of flagellar genes was induced by adding anhydrotetracycline. Following induction, 100μl samples were collected every 10 mins. Cultures were briefly pelleted, resuspended into buffered saline and kept on ice. For microscopy, 5 µl of the cell suspension was placed on a large coverslip and overlaid with a 1% agarose pad (53). A small cover slip was placed on top of the agarose pad to prevent drying during imaging. Samples were imaged using a Zeiss Axio Observer inverted microscope equipped with Pln Apo 100x/1.4 Oil immersion objective (Ph3), YFP, CFP and mCherry filters, an Axiocam 506 mono CCD-camera and Colibri 7 LED light source (Zeiss). Images were analyzed using the Intellesis trainable object classification in the Zen 3.12 software. Three independent cultures were analyzed for each genetic background and/or time point. For Figure 2, the total number of cells expressing *cfp* (class 3) was counted relative to the total number of cells expressing *yfp* (class 2 in different genetic backgrounds. For the time course experiment, the number of cells expressing class 2 (YFP) or class 3 (CFP) was counted, relative to the total number of cells (expressing constitutive mCherry). The time-course figure was plotted using Prism 10 software.

### Motility Assays

Single colonies from freshly streaked bacteria were inoculated into soft motility agar (10 g/l tryptone, 5 g/l NaCl, 3g/l Difco-Bacto-Agar) by stabbing the plate. The diameter of swimming halo diameters was measured after incubation at 37 °C for 6 h, and values were compared to the parent strain. At least 3 colony replicates samples were performed per sample.

### Minimal Inhibitory Concentration (MIC) assays

Assessment of early and late substrates secretion was performed in different genetic backgrounds using β-lactamase (lacking its Sec-dependent signal sequence) fused to either the C-terminus of FlgE (as described in (36)), FliK or FlgM (9). FlgE-Bla, FliK-Bla and FlgM-Bla secretion into the periplasm was evaluated using minimal inhibitory concentration (MIC) assays as described (10). The procedure was conducted as follows: a series of LB-ampicillin (Ap) solutions was prepared using 2-fold serial dilutions starting from a freshly made LB solution containing 800 µg/ml Ap. A total of 198 µl of each dilution was dispensed into wells of a clear 96-well plate. Independent bacterial cultures were grown overnight in 1 ml of LB medium with aeration at 37 °C. These overnight cultures were diluted 200-fold in PBS, and 2 µl of the diluted cultures was added to each well containing the LB-Ap solutions. Plates were incubated at 37 °C for 18 hours in the dark with shaking at 180 rpm. The minimum inhibitory concentration (MIC) was defined as the lowest Ap concentration at which no visible bacterial growth occurred. Each strain was tested in at least three independent biological replicates.

### FliK dependent release of the cleavable C-terminal domain of FlhB

Strain TH27122 (Δ*araBAD957*::*rflP⁺* Δ*rflM8403 flhDC5451*::TPOP *fljB^enx^ vh2*) was grown overnight in LB broth at 37 °C. The next day, the culture was subcultured into four flasks containing 250 ml of LB medium and grown at 37 °C to an OD₆₀₀ of 0.5. At that point, anhydrotetracycline (ATc) was added to induce the flagellar system. Cells were harvested by centrifugation (5,000 × g, 30 min, 4 °C) at 20, 30, 40, and 50 min after ATc addition, respectively. Cell pellets were resuspended in 10 ml of ice-cold membrane buffer (50 mM Tris·HCl, pH 7.5, 50 mM NaCl, 250 mM sucrose, 0.5 mM DTT, 10% glycerol, and 2 mM CaCl₂) and disrupted by passage through a French press. DNase I (10 µg/ml) and MgCl₂ (5 mM) were added, and the lysates were incubated for 10 min at 22 °C. Unbroken cells were removed by low-speed centrifugation (5,000 × g, 10 min, 4 °C), and membranes were collected by ultracentrifugation (100,000 × g, 60 min, 4 °C). Membrane pellets were resuspended in SDS/PAGE sample buffer and analyzed by electrophoresis on 16.5% SDS/PAGE gels, followed by Western blotting using anti-FlhB C-terminal antibodies (see below).

Strains TH27622 (*flhDC5451*::TPOP Δ*rflM8403 fljB^enx^ vh2*) and TH27623 (*flhDC5451*::TPOP Δ*rflM8403* Δ*fliK6140 fljB^enx^ vh2*) were subcultured from overnight cultures into 50 ml of LB medium and grown at 37 °C to an OD₆₀₀ of 0.5. ATc was added and cultures were incubated for 30 min before cells were collected by centrifugation, washed once with buffered saline, and resuspended in fresh LB medium. Cultures were incubated for an additional 30 min in LB media without ATc, harvested, and analyzed by SDS/PAGE and Western blotting as above using anti-FlhB C-terminal antibodies.

### Secretion assays

Strains TH30093, TH30110 and TH30111 were grown overnight in LB broth at 37 °C. The following day, 10 ml of the overnight culture was sub-cultured into 1L of defined minimal E-medium containing 0.2% glycerol and amino acid pools 1–5, 8, and 11 (described on pages 207-209 in (14)) and grown to an OD₆₀₀ of 0.5. Cells were pelleted and resuspended in 20 ml of defined medium supplemented with 0.2% arabinose to induce expression of *fliK* or *fliK–TOP7*. After 15 min of shaking, a 1 ml aliquot was pelleted and resuspended in 200 µl of SDS/PAGE sample buffer for western blot analysis. The remaining culture was pelleted, and the supernatant was filtered through a 0.2 µm PES membrane and concentrated to 200 µl using an Amicon Ultra-4 centrifugal filter unit with a 3 kDa molecular weight cutoff (Millipore).

### Western blots

Samples diluted in SDS/PAGE sample buffer were heated for 5 min at 95 °C and separated on 4-20% Mini-PROTEAN TGX gels (Bio-Rad) or 16.5% acrylamide SDS-PAGE gels. Proteins were transferred to Trans-Blot Turbo Mini LF PVDF membranes using the Trans-Blot Turbo transfer system (Bio-Rad). Membranes were probed with polyclonal affinity-purified rabbit antibodies, including anti-FliK (raised against full protein), anti-FlgE (raised against full protein), anti-FlgM (raised against amino acids 16–34, QTRETSDTPVQKTRQEKTS; LI International) and anti–C-terminal FlhB (raised against amino acids 354–383, RLAGGQRPPQPENLPVPEALD FMNEKNTDG; LI International). IRDye 800CW Goat anti-rabbit secondary antibodies (LI-COR) were used for detection. Signals were visualized by infrared fluorescence using a Sapphire FL Biomolecular Imager (Azure Biosystems).

### Assay for the timing of class-3 Specificity Switching

Class 3 promoter activity was monitored over time using a chromosomally integrated P*_fliC_*-*luxCDABE* reporter (from *Photorhabdus luminescens*) (20). Strains TH30413 (*fliK*-Bla, WT), TH30414 (*fliK* (Δaa311–320)-Bla), and TH301415 (*fliK* (P296L)-Bla) each carried a tetracycline-inducible *flhDC* promoter to initiate flagellar gene expression, along with the *P_fliC_-luxCDABE reporter.* Cultures (2 mL) were grown at 37 °C in LB medium to an optical density (OD₆₀₀) of 0.5, at which point anhydrotetracycline was added to a final concentration of 125 ng/µL. Cells (200 µL) were transferred to a 96-well plate and incubated at room temperature in a PolarStar Optima plate reader. Luminescence was measured every 5 min with intermittent shaking. For each strain, four technical replicates were analyzed for each of three biological replicates.

### Bacterial two-hybrid

The bacterial two-hybrid (BACTH) system, based on the reconstitution of adenylate cyclase activity in an *Escherichia coli cya* mutant (25), was used to test interactions between FlhA and either Flk or FliK. The cytoplasmic domain of FlhA (residues S362–K692) was cloned into the pKNT25 vector using *XbaI* and *KpnI*. Full-length *fliK* and *flk* were cloned into the pUT18C vector using *XbaI* and *KpnI*, and *SalI* and *BamHI*, respectively. Plasmid combinations, including empty-vector controls, were electroporated into *E. coli* BTH101 cells. After recovery in LB medium for 30 min at 37 °C, cells were plated on LB agar containing ampicillin and kanamycin and incubated overnight at 37 °C. Transformants were cultured for 4 h in liquid LB with antibiotics and then spotted onto MacConkey-lactose agar supplemented with ampicillin and kanamycin. Plates were incubated at 30 °C for 2 days. Red (Mac-Lac⁺) colonies indicated a positive interaction, whereas white (Mac-Lac⁻) colonies indicated no interaction.

### Structural modeling and visualization

Structural models of the C-terminal domains of FliK (residues 269–end) and the cleaved form of FlhB (residues 270–end) were generated using the AlphaFold3 prediction server (1). The closed (early secretion) and open (late secretion) forms of the FlhA_C_ nonameric ring were constructed using the crystal structures with PDB IDs 3MYD and 3A5I, respectively. The D1 domain from each crystal structure was superimposed onto the corresponding domain of the 7AMY structure using UCSF ChimeraX (version 1.10.1) (46).

## Acknowledgements

This work was supported by PHS grants R01GM056141 and R56AI184395 from the National Institutes of Health (to K.T.H.) and by funding from the Undergraduate Research Opportunities Program at the University of Utah (awarded to DB. Wu and C.T. Mellor). D. Niketic was funded by Genetics Training Grant T32-GM007464 from the National Institutes of Health. This work was also supported by JSP KAKENHI Grant Number JP19H03182, JP22H02573, and JP22K19274 (to T.M.). MEXT KAKENHI Grant Number JP20H05532 and JP22H04844 (to T.M.), by Research Support Project for Life Science and Drug Discovery (BINDS) from AMED under Grant Number JP25ama121003 (to K.N.), and by JEOL YOKOGUSHI Research Alliance Laboratories of The University of Osaka (to K.N). We thank Paige Wheatley for help with the bacterial two-hybrid assay.

## Notes

### Competing Interest Statement

The authors have declared no competing interest.

