## Supplemental for "Active removal of inhibitory components drives the flagellar Type III Secretion Specificity Switch"

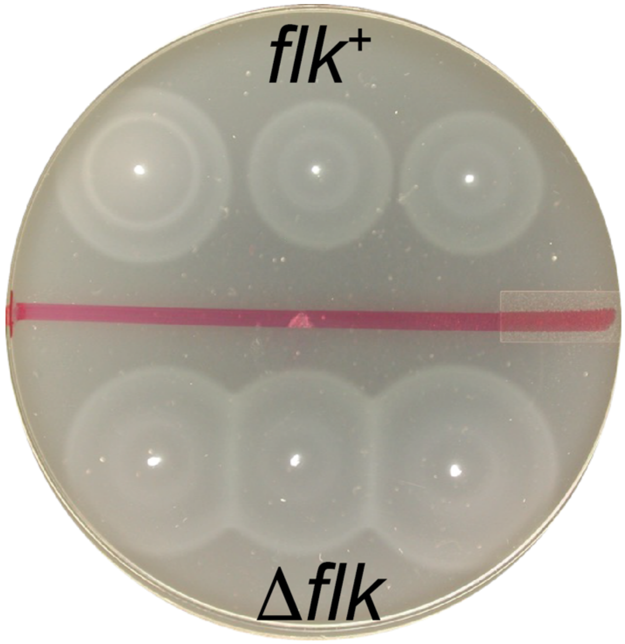

**Supplemental Figure 1.** The deletion of *flk* has no apparent effect on *Salmonella* motility (*flk*$^+$ = strain TH437 (LT2); Δ*flk* = strain TH1460).

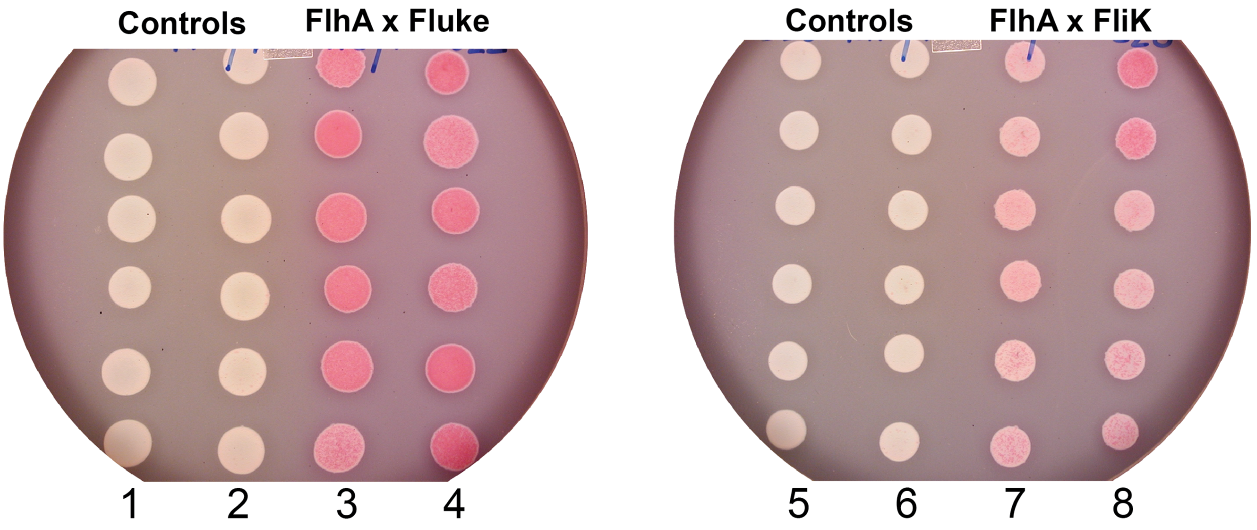

**Supplemental Figure 2.** A bacterial two hybrid screen for interaction between either Fluke or FliK with $FlhA_C$ indicates a positive interaction phenotype ($Lac^+$ pink color formation). Lane 1= pUT18C-FliK x pKNT25 empty vector; lane 2= pUT18C empty vector x pKNT25-FlhA C-terminus; lanes 3 and 4 = pUT18C-Fluke x pKNT25-FlhA C-terminus; lane 5= pUT18C-Fluke x pKNT25 empty vector; lane 6= pUT18C empty vector x pKNT25-FlhA C-terminus; lanes 7 and 8 = pUT18C-FliK x pKNT25-FlhA C-terminus.

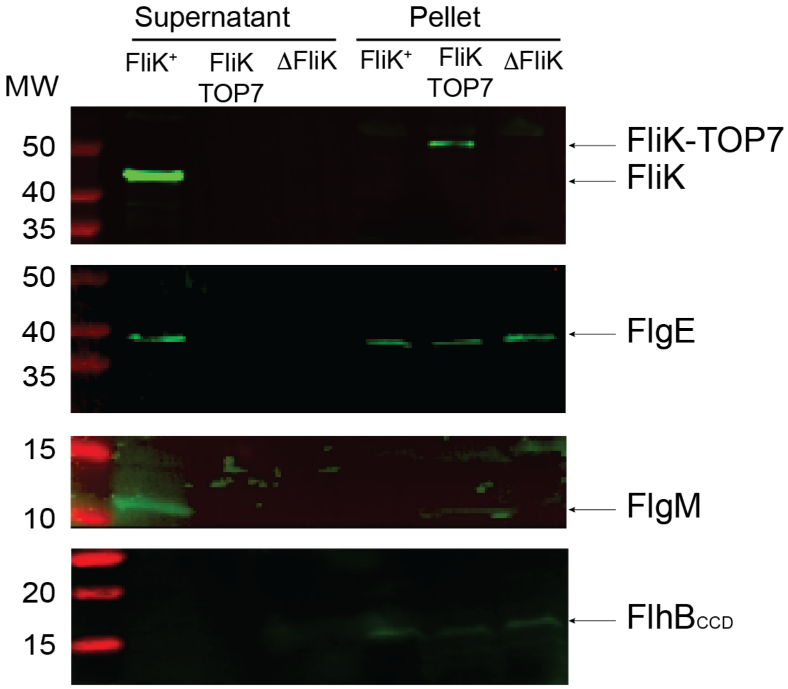

**Supplemental Figure 3. The cleavable C-terminal domain of FlhB is not detected in the cell supernatant, just after the secretion specificity switch has occurred.** Cells (1L) were grown to OD 0.5. Cells were pelleted, washed once and resuspended into 20 ml of media containing arabinose and induced for 15 mins. Supernatant was concentrated to 200 µl and 20 µl/lane was loaded.

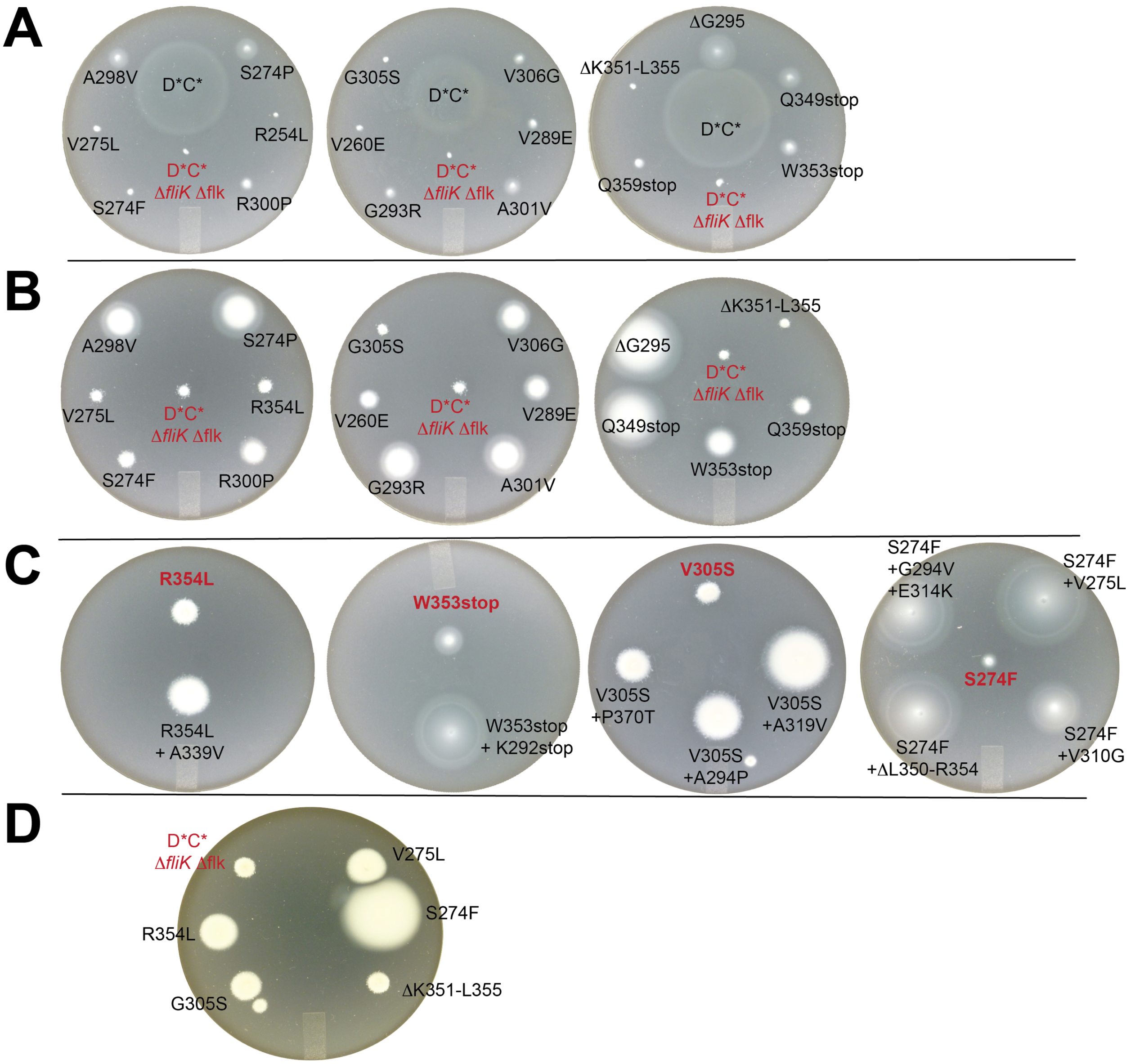

**Supplemental Figure 4. Motile revertants of Δ*fliK* Δ*flk* double mutant strain in the *flhD*C** background TH29848 (*flhD8070*(L22H) *flhC8092*(Q29P) Δ*fliK9249* Δ*flk-7755*). A.** Motility relative to the parent TH29848 (labeled red) and a *fliK*$^{+}$ *flk*$^{+}$ motile strain showing that motile revertants do not exhibit full "wild-type" motility after 4 hours incubation in soft agar motility plates at 37°C. **B.** Motility phenotypes relative to the parent strain (red), TH29848, after 8 hours incubation in soft agar medium at 37°C. **C.** Double-mutants with increased motility shown relative to the original motile revertant phenotype (red) after 10 hours incubation in soft agar motility plates at 37°C. **D.** These are motility revertants with only slight increased motility phenotypes after 16 hours incubation on soft agar motility plates at 37°C.

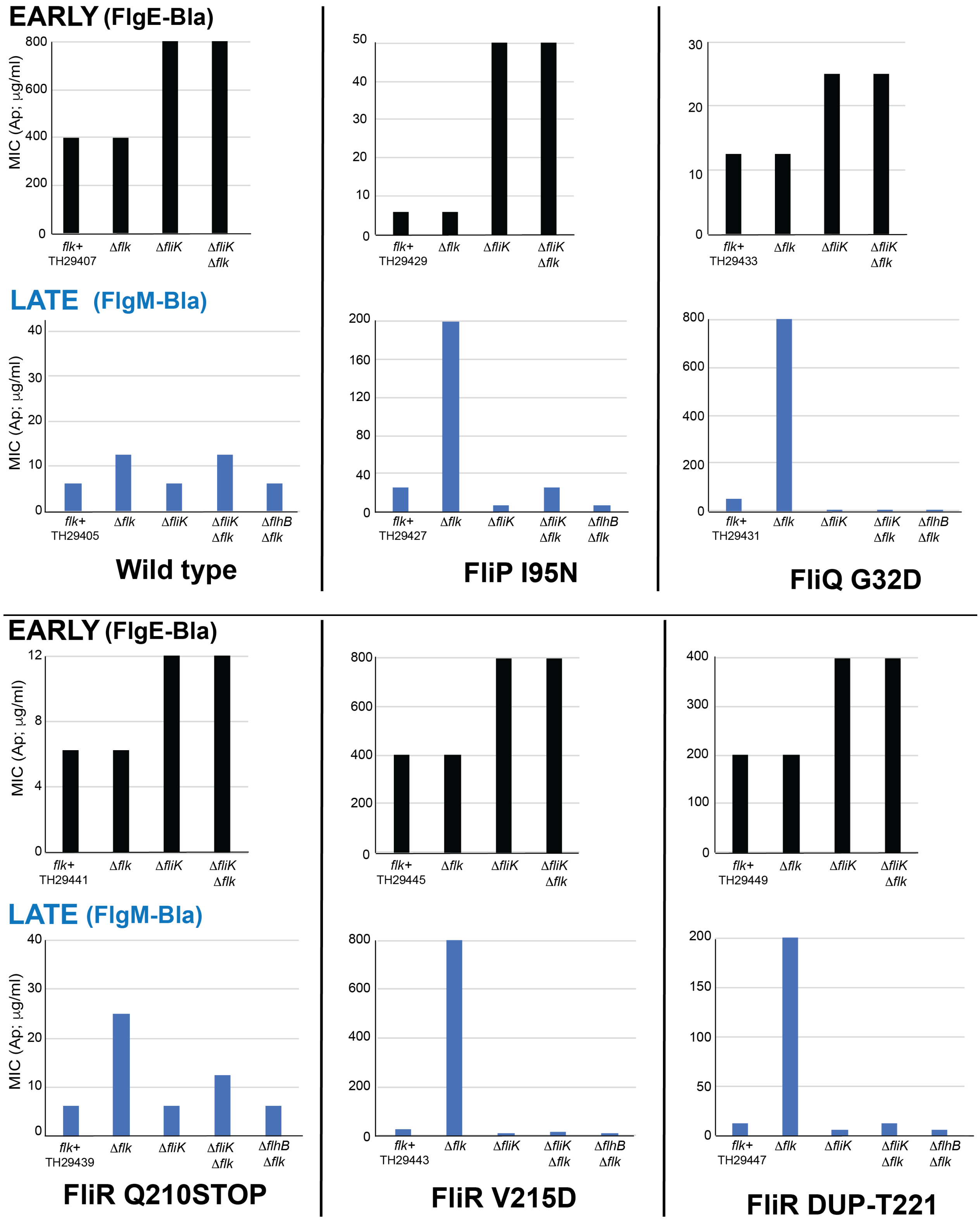

**Supplemental Figure 5.** Levels of early (FlgE-Bla) and late (FlgM-Bla) substrate secretion in the FlhB-bypass mutants in the presence and absence of Fluke, and/or FliK. Parent strain numbers (*flk*$^{+}$) are indicated in the figure. Genotypes of all strains used are listed in Table S4.

**Supplemental Tables**

Table S1. Plasmids and Bacterial strains used in this study*

**Plasmids:**

| Plasmid Name | Usage and relevant characteristic | Source* |
|---|---|---|
| pKD3 | For inserting FRT-Cm-FRT chromosomal cassette, $Ap^R$ plasmid | (13) |
| p-Sim5 | Temperature inducible λ-Red expressing plasmid, $Cm^R$ plasmid, pSC101 origin of replication (replicates at 30°C) | (14) |
| p-Sim-6 | Temperature inducible λ-Red expressing plasmid, $Ap^R$ plasmid, pSC101 origin of replication (replicates at 30°C) | (14) |
| p-EM5 | $Km^R$ $P_{nahG}$-*fliQ*$^+$ | M. Erhardt |

***E.coli* strains:**

| ***E.coli* Strains** | Genotype | Source* |
|---|---|---|
| MG1655 MGRFC | Δ*leuB motA*(E98K) Int5::$Zeo^R$-PRNA-mcherry *galK*::$P_{fliF}$(*E. coli*)-CFP-$Ap^R$ *attB*::$P_{fliC}$(*E. coli*)-YFP-$Km^R$ | (33) |
| TH408 | *nad*::Tn*10d*Tc | Lab collection |
| BTH101 | $F^-$ *cya-99 araD139 galE15 galK16 rpsL1*($Str^R$) *hsdR2 mcrA1 mcrB1* | BACTH, Euromedex |

***Salmonella* Typhimurium strains:**

| *Salmonella* strains | Genotype | Source* |
|---|---|---|
| TH437 | LT2 *Salmonella* wild type strain | J.Roth |
| TH9949 | *flg*E*6569*::*bla* Δ*flgBC6557* | Lab collection |
| TH13359 | *fliK7582*::*bla* | Lab collection |
| TH13807 | *fliK7582*::*bla* Δ*flgA7656* (Δamino acids 6-214 (of 219)) | Lab collection |
| TH13808 | *fliK7582*::*bla* Δ*flgB7657* (Δamino acids 6-133(of 138)) | Lab collection |
| TH13809 | *fliK7582*::*bla* Δ*flgC7658* (Δamino acids 6-129(of 134)) | Lab collection |
| TH13810 | *fliK7582*::*bla* Δ*flgD6540* (Δamino acids 2-220(of 232)) | Lab collection |
| TH13811 | *fliK7582*::*bla* Δ*flgE7659* (Δamino acids 6-398(of 403)) | Lab collection |
| TH13812 | *fliK7582*::*bla* Δ*flgF7660* (Δamino acids 6-246(of 251)) | Lab collection |
| TH13813 | *fliK7582*::*bla* Δ*flgG7661* (Δamino acids 6-255(of 260)) | Lab collection |
| TH13814 | *fliK7582*::*bla* Δ*flgH7662* (Δamino acids 6-227(of 232)) | Lab collection |
| TH13815 | *fliK7582*::*bla* Δ*flgI7663* (Δamino acids 6-360(of 365)) | Lab collection |
| TH13816 | *fliK7582*::*bla* Δ*flgJ7664* (Δamino acids 6-311(of 316)) | Lab collection |
| TH13817 | *fliK7582::bla* Δ*flgK7665* (Δamino acids 6-548(of 553)) | Lab collection |
| TH13818 | *fliK7582*::*bla* Δ*flgL7666* (Δamino acids 6-312(of 317)) | Lab collection |
| TH14607 | Δ*flk-7755* | Lab collection |
| TH23959 | Δ*flhB8612*$_{\Delta CCD}$ (*flhB*$_{\Delta CCD}$ = deletion of amino acids P270-G383) | Lab collection |
| TH24321 | Δ*flgBC6557 flgE6569*::*bla* Δ*flk-7755* | |
| TH24658 | Δ*flgBC6557 flgE6569*::*bla* Δ*flk-7755* | |
| TH24722 | *flgM6427::bla* Δ*flgB-L8735* Δ*flhB8612*$_{\Delta CCD}$ *flhD8070 flhC8092 fliA5225*(H14D) Δ*fliB-T7771 fljB*$^{enx}$ *vh2* | |
| TH24787 | *flgM6427::bla* Δ*flgB-L8735* Δ*flhB8612*$_{\Delta CCD}$ *flhD8070 flhC8092 fliA5225*(H14D) Δ*fliB-T7771* Δ*flk-7755 fljB*$^{enx}$ *vh2* | |
| Salmonella strains | Genotype | Source* |
| TH24789 | Δ*araBAD961*::*flk*$^{+}$ *flgM6427::bla* Δ*flgB-L8735* Δ*flhB8612*$_{\Delta CCD}$ *flhD8070 flhC8092 fliA5225*(H14D) Δ*fliB-T7771 fljB*$^{enx}$ *vh2* | |
| TH24800 | *flgM6427*::*bla* Δ*flgB-L8735* Δ*flhB8612*$_{\Delta CCD}$ *flhD8070 flhC8092 fliA5225*(H14D) Δ*fliB-T7771* Δ*fliK6140* Δ*flk-7755 fljB*$^{enx}$ *vh2* | |
| TH24819 | *flgM6427*::*bla* Δ*flgB-L8735* Δ*flhB8612*$_{\Delta CCD}$ *flhD8070 flhC8092 fliA5225*(H14D) Δ*fliB-T7771* Δ*fliK6140 fljB*$^{enx}$ *vh2* | |

| | | |
|---|---|---|
| TH25226 | *flgM6427*::*bla* Δ*flgB-L8735 flhD8070 flhC8092 fliA5225*(H14D) Δ*fliB-T7771* Δ*flk-7755* $fljB^{enx}$ *vh2* | |
| TH25935 | *flgM6427*::*bla* ΔflgB-L8735 $flhB8612_{\Delta CCD}$ $fljB^{enx}$ *vh2* | |
| TH25936 | Δ*flgBC6557 flgE6569*::*bla* $flhB861_{\Delta CCD}$ *2* $fljB^{enx}$ *vh2* | |
| TH25938 | *flgM6427*::*bla* Δ*flgB-L8735* $flhB8612_{\Delta CCD}$ Δ*flk-7755* $fljB^{enx}$ *vh2* | |
| TH25939 | Δ*flgBC6557 flgE6569*::*bla* $flhB8612_{\Delta CCD}$ Δ*flk-7755* $fljB^{enx}$ vh2 | |
| TH26028 | *flgM6427*::*bla* Δ*flgB-L8735* $flhB8612_{\Delta CCD}$ *fliK6620* Δ*flk-7755* $fljB^{enx}$ *vh2* | |
| TH26992 | *flgM6427*::*bla* Δ*flgB-L8735 STM1911*::Tn*10d*Tc Δ*flhBAE7670*::FCF *flhD8070 flhC8092 fliA5225*(H14D) Δ*fliB-T7771* Δ*flk-7755* $fljB^{enx}$ *vh2* | |
| TH27083 | pSIM5/LT2 | |
| TH27122 | Δ*araBAD957*::$rflP^+$ Δ*rflM8403* $P_{flhDC}$*5451*::TPOP $fljB^{enx}$ *vh2* | |
| TH27152 | *flgM6427*::*bla* Δ*flgB-L8735 flhD8070 flhC8092 fliA5225*(H14D) Δ*fliB-T7771* Δ*fliQ8223*::*tetRA* (Δ codons 31-50) Δflk-7755 $fljB^{enx}$ *vh2* | |
| TH27337 | *attB*::$Cm^R$-$P_{fliF}$-YFP *argW*::$Zeo^R$-mcherry Δ*galK*::$Ap^R$-$P_{fliC}$-CFP $P_{flhDC}$*5451*::TPOP $fljB^{enx}$ *vh2* | |
| TH27343 | *attB*::$Cm^R$-$P_{fliF}$-YFP *argW*::$Zeo^R$-mcherry Δ*galK*::$Ap^R$-$P_{fliC}$-CFP *flgM7929*::TPOP $fljB^{enx}$ *vh2* | |
| TH27622 | $P_{flhDC}$*5451*::TPOP Δ*rflM8403* $fljB^{enx}$ *vh2* | |
| TH27623 | $P_{flhDC}$*5451*::TPOP Δ*rflM8403* Δ*fliK6140* $fljB^{enx}$ *vh2* | |
| TH27965 | Δ*araBAD2101*::$flhB^+$ *flgM6427*::*bla* Δ*flgB-L8735 flhD8070 flhC8092 fliA5225*(H14D) Δ*fliB-T7771* Δ*flk-7755* $fljB^{enx}$ *vh2* | |
| TH28049 | *attB*::$Cm^R$-$P_{fliF}$-YFP Δ*galK*::$Ap^R$-$P_{fliC}$-CFP $fljB^{enx}$ *vh2* | |
| TH28050 | *attB*::$Cm^R$-$P_{fliF}$-YFP Δ*galK*::$Ap^R$-$P_{fliC}$-CFP Δ*flgHI958* $fljB^{enx}$ *vh2* | |
| TH28051 | *attB*::$Cm^R$-$P_{fliF}$-YFP Δ*galK*::$Ap^R$-$P_{fliC}$-CFP *flhB8612* $fljB^{enx}$ *vh2* | |
| TH28052 | *attB*::$Cm^R$-$P_{fliF}$-YFP Δ*galK*::$Ap^R$-$P_{fliC}$-CFP Δ*flk-7755* $fljB^{enx}$ *vh2* | |
| TH28053 | *attB*::$Cm^R$-$P_{fliF}$-YFP Δ*galK*::$Ap^R$-$P_{fliC}$-CFP Δ*flgHI958* Δ*flk-7755* $fljB^{enx}$ *vh2* | |
| TH28054 | *attB*::$Cm^R$-$P_{fliF}$-YFP Δ*galK*::$Ap^R$-$P_{fliC}$-CFP $flhB8612_{\Delta CCD}$ Δ*flk-7755* $fljB^{enx}$ *vh2* | |
| TH28055 | *attB*::$Cm^R$-$P_{fliF}$-YFP Δ*galK*::$Ap^R$-$P_{fliC}$-CFP $flhB8612_{\Delta CCD}$ Δ*flgHI958* $fljB^{enx}$ *vh2* | |
| TH28056 | *attB*::$Cm^R$-$P_{fliF}$-YFP Δ*galK*::$Ap^R$-$P_{fliC}$-CFP $flhB8612_{\Delta CCD}$ Δ*flgHI958* Δ*flk-7755* $fljB^{enx}$ *vh2* | |
| TH28157 | *flgM6427*::*bla* Δ*flgB-L8735* $fljB^{enx}$ *vh2* | |
| TH28158 | *flgM6427*::*bla* Δ*flgB-L8735* Δ*flk-7755* $fljB^{enx}$ *vh2* | |
| TH29063 | Δ*araBAD2101*::$flhB^+$ *flgM6427*::*bla* Δ*flgB-L8735* $flhB8612_{\Delta CCD}$ Δ*flk-7755* $fljB^{enx}$ *vh2* | |
| TH29064 | Δ*araBAD2101*::$flhB^+$ Δ*flgBC6557 flgE6569*::*bla* $flhB8612_{\Delta CCD}$ Δ*flk-7755* $fljB^{enx}$ *vh2* | |
| TH29116 | Δ*flgBC6557 flgE6569*::*bla* Δ$flhB8612_{\Delta CCD}$ Δ*fliK9249* Δ*flk-7755* | |
| TH29314 | *attB*::$Cm^R$-$P_{fliF}$-YFP Δ*galK*::$Ap^R$-$P_{fliC}$-CFP Δ*flk-7755* $fljB^{enx}$ *vh2* | |
| TH29315 | *attB*::$Cm^R$-$P_{fliF}$-YFP Δ*galK*::$Ap^R$-$P_{fliC}$-CFP *flhB7152*(N269A) Δ*flgHI958* $fljB^{enx}$ *vh2* | |
| TH29316 | *attB*::$Cm^R$-$P_{fliF}$-YFP Δ*galK*::$Ap^R$-$P_{fliC}$-CFP *flhB7152*(N269A) Δ*flgHI958* Δ*flk-7755* $fljB^{enx}$ *vh2* | |
| TH29317 | *attB*::$Cm^R$-$P_{fliF}$-YFP Δ*galK*::$Ap^R$-$P_{fliC}$-CFP $flhB8612_{\Delta CCD}$ *flhD8070 flhC8092* Δ*flk-7755* $fljB^{enx}$ *vh2* | |
| TH29318 | *attB*::$Cm^R$-$P_{fliF}$-YFP Δ*galK*::$Ap^R$-$P_{fliC}$-CFP $flhB8612_{\Delta CCD}$ *flhD8070 flhC8092* Δ*flgHI958* $fljB^{enx}$ *vh2* | |
| *Salmonella* strains | Genotype | Source* |

| | |
|---|---|
| TH29319 | *attB*::Cm$^{R}$-P$_{fliF}$-YFP Δ*galK*::Ap$^{R}$-P$_{fliC}$-CFP *flhB8612*$_{\Delta CCD}$ *flhD8070 flhC8092* Δ*flgHI958* Δ*flk-7755 fljB*$^{enx}$ *vh2* |
| TH29357 | *attB*::Cm$^{R}$-P$_{fliF}$-YFP Δ*galK*::Ap$^{R}$-P$_{fliC}$-CFP Δ*flhB8789 (*leaves first 5 and last 9 amino acids of FlhB) *flhD8070 flhC8092* Δ*flk-7755 fljB*$^{enx}$ *vh2* |
| TH29358 | *attB*::Cm$^{R}$-P$_{fliF}$-YFP Δ*galK*::Ap$^{R}$-P$_{fliC}$-CFP Δ*flgHI958 flhB8789 flhD8070 flhC8092 fljB*$^{enx}$ *vh2* |
| TH29359 | *attB*::Cm$^{R}$-P$_{fliF}$-YFP Δ*galK*::Ap$^{R}$-P$_{fliC}$-CFP Δ*flgHI958 flhB8789 flhD8070 flhC8092* Δ*flk-7755 fljB*$^{enx}$ *vh2* |
| TH29848 | *flhD8070 flhC8092* Δ*fliK9249* Δ*flk-7755* |
| TH30093 | Δ*ara*BAD7606::*fliK*$^{+}$ Δ*fliK9249 attB*::Cm$^{R}$-P$_{fliF}$-YFP Δ*galK*::Ap$^{R}$-P$_{fliC}$-CFP *flhD8070 flhC8092* Δ*prgH73*::*tetRA ompT*::Km *fljB*$^{enx}$ *vh2* |
| TH30110 | Δ*araBAD7609*::*fliK*-TOP7 Δ*fli*K9249 *attB*::Cm$^{R}$-P$_{fliF}$-YFP Δ*galK*::Ap$^{R}$-P$_{fliC}$-CFP *flhD8070 flhC8092* Δ*prgH73*::*tetRA ompT*::Km *fljB*$^{enx}$ *vh2* |
| TH30111 | Δ*araBAD937*::FKF Δ*fliK9249 attB*::Cm$^{R}$-P$_{fliF}$-YFP Δ*galK*::Ap$^{R}$-P$_{fliC}$-CFP *flhD8070 flhC8092* Δ*prgH73*::*tetRA ompT*::Km *fljB*$^{enx}$ *vh2* |
| TH30250 | Δ*flgBC6557 flgE6569*::*bla* Δ*flhB8612*$_{\Delta CCD}$ Δ*fliK9249* |
| TH30367 | *fliK9504*(Δaa311-320) *fliK7582*::*bla* |
| TH30368 | *fliK9505*(P296L) *fliK7582*::*bla* |
| TH30413 | Δ*galK*::P$_{fliC}$-*luxCDBAE*-Cm$^{R}$ P$_{flhDC}$*5451*::Tn*10d*Tc[*del-25*] *fliK7582*::*bla* |
| TH30414 | Δ*galK*::P$_{fliC}$-*luxCDBAE*-Cm$^{R}$ P$_{flhDC}$*5451*::Tn*10d*Tc[*del-25*] *fliK9504*(Δaa311-320) *fliK7582*::*bla* |
| TH30415 | Δ*galK*::P$_{fliC}$-*luxCDBAE*-Cm$^{R}$ P$_{flhDC}$*5451*::Tn*10d*Tc[*del-25*] *fliK9505*(P296L) *fliK7582*::*bla* |

*Unless indicated otherwise these strains were constructed during this work

### ***Salmonella enterica serovar Typhimurium*** **14028 strains:**

| Salmonella strains | Genotype | Source* |
|---|---|---|
| TH27356 | 14028 *argW*::Zeo$^{R}$-PmCherry *attB*::Km$^{R}$-P$_{fliF}$-YFP Δ*galK*::Ap$^{R}$-P$_{fliC}$-CFP Δ*prgH74* Δ*ssaN109* Δ*flgHI958 motA8739*(E98K) Δ*fliC7716 fljB*$^{enx}$ *vh2* | |
| TH27359 | 14028 *argW*::Zeo$^{R}$-PmCherry *attB*::Km$^{R}$-P$_{fliF}$-YFP Δ*galK*::Ap$^{R}$-P$_{fliC}$-CFP Δ*prgH74* Δ*ssaN109* Δ*flgHI958* Δ*flk*-7755 *motA8739*(E98K) Δ*fliC7716 fljB*$^{enx}$ *vh2* | |
| TH27476 | 14028 *argW*::Zeo$^{R}$-PmCherry *attB*::Km$^{R}$-P$_{fliF}$-YFP Δ*galK*::Ap$^{R}$-P$_{fliC}$-CFP Δ*prgH74* Δ*ssaN109* Δ*flgHI958* Δ*flk-7755 flhB8612*$_{\Delta CCD}$ *motA8739* Δ*fliC7716 fljB*$^{enx}$ *vh2* | |
| TH27477 | 14028 *argW*::Zeo$^{R}$-PmCherry *attB*::Km$^{R}$-P$_{fliF}$-YFP Δ*galK*::Ap$^{R}$-P$_{fliC}$-CFP Δ*prgH74* Δ*ssaN109* Δ*flgHI958 flhB8612 motA8739* Δ*fliC7716 fljB*$^{enx}$ *vh2* | |
| TH27478 | 14028 *argW*::Zeo$^{R}$-PmCherry *attB*::Km$^{R}$-P$_{fliF}$-YFP Δ*galK*::Ap$^{R}$-P$_{fliC}$-CFP Δ*prgH74* Δ*ssaN109* Δ*flgHI958* Δ*flk-7755* Δ*fliK6620 motA8739* Δ*fliC7716 fljB*$^{enx}$ *vh2* | |
| TH27451 | 14028 *argW*::Zeo$^{R}$-PmCherry *attB*::Km$^{R}$-P$_{fliF}$-YFP Δ*galK*::Ap$^{R}$-P$_{fliC}$-CFP Δ*prgH74* Δ*ssaN109* Δ*flgM5628*::FRT Δ*flgHI958 motA8739* Δ*fliC7716 fljB*$^{enx}$ *vh2* | |

Table S2. Oligonucleotides used in this study

| Name | Sequence (from 5'- to 3'-) |
|---|---|
| 1874-flhBCLtetR | gatggaagatgtgccgaaagcggacgtcattgtcactaacttaagacccactttcacatt |
| 1910-flhB375tetA | cgcgaccagattagccatcagtattcttctcgttcataaactaagcacttgtctcctg |
| 1911-flhBccDel-A | gcgcagcgccgcatgatggaagatgtgccgaaagcggacgtcattgtcactaactaattt |
| 1912-flhBccDel-B | cgcagcatcgcgaccagattagccatcagtattcttctcgttcataaattagttagtgac |
| 4543-fliKfullbla_rv | gatcttcagcatcttttactttcaccagcgtttctgggtgggcgaagatatccactgcgc |
| 7516-FliK278 | gtcatgttatttacgcgtcag |
| 8283-galK-sCFP3A | cgcggtcagcgacatccattttcgcgaatccggagtaTAAaaAGGTCTAGGCGGCGCCTA |
| 8284-galK-amp-cfp3A | cctgctccttgtgacggtttgcatacataaaaggtttcTTACCAATGCTTAATCAGTGAGGC |
| 8285-ybhCvenusNB | gcctgaaaaggaactttttaccttttcgccttcccgtttcgtGGCAGCAAAACCCGTACC |
| 8286-ybhC venusNBKan | agttaaatgacatccattgaagcctgctttttttatactaagttgaGCTTGGATTCTCACC |
| 8309-attB-Km_Prom-venusNB-tetR | acctataaaaataggcgtatcacgaggccctttcgtcttgac TTAAGACCCACTTTCACATT |
| 8315-galKtetR-scCFP3A | atttgaatgtatttagaaaaataaacaaataggggttccgcgTTAAGACCCACTTTCACATT |
| 8331-galK-tetA-CFP | ggtctagactccttactaaagttaaacaaaattattatcaatCTAAGCACTTGTCTCCTG |
| 8458-galK-PFliCsal | atttgaatgtatttagaaaaataaacaaataggggttccgcgGTTCTTTGTCAGGTCTGTC |
| 8459-attB-Km_PfliF-salm-venusfw | actcatatgtatatctccttcttaaagttaaacaaaattatt GGATTCGCGCGTAGGCGA |
| 8460-attB-Km_PfliFsal-venusrv | cctataaaaataggcgtatcacgaggccctttcgtcttgac CGTCGACTGCGAGTGC |
| 9083-new-attB-km-promvenusNBtetA | actcatatgtatatctccttcttaaagttaaacaaaattattCTAAGCACTTGTCTCCTG |
| 9142-galK-PFliC-RBSforCFP | ggtctagactccttactaaagttaaacaaaattattatcaatCGCAGACCGGAAGACAGA |
| 9305-attB-Cm-rv | tgagcttggattctcaccaataaaaaacgcccggcggcaaccCACTCATCGCAGTACTGTTGTAT |
| 9306-attB-Cm-fw | tttcttagacgtcggaattgccagctggggcgccctctggTCCTGGTGTCCCTGTTGATac |
| 9318-FliQtetR | tctcgccctggctgcgccgctgttactcgtcgcgctgattttaagacccactttcacatt |
| 9319-FliQtetA | aacgataattgcgatgaataccgcgacgattttagggataaactaagcacttgtctcctg |
| 9320-FliQ-AA31-50dopped | gttactcgtcgcgctgattaccggcctcattatcagcatcttgcaggccgcgactcagattaatgaaatgacgctgtcgtttatc cctaaaatcgtcgcg |
| 9321-fliQ50-fill | cgcgacgattttagggat |
| 9322-FliQ-30fw | agtcgctctcgccctggctgcgccgctgttactcgtcgcgctgat |
| 9323-FliQ-50-rev | ggcaacgataattgcgatgaataccgcgacgattttagggat |

**Table S3. Class 3 promoter activity relative to class 2 promoter activity in individual bacterial cells, in different genetic backgrounds**

| Strain Number | Relevant Genotype (all strains contained the $P_{fliF}$-*yfp* $P_{fliC}$-*cfp*) | Number of cells expressing YFP* | Number of cells expressing CFP* | Percentage Class3/2 |
|---|---|---|---|---|
| TH28049 | wild type | 992 | 989 | 100 |
| TH27343 | *flgM* null | 853 | 854 | 100 |
| TH28052 | Δ*flk* | 874 | 866 | 99 |
| TH28050 | Δ*flgHI* | 1016 | 1 | <0.01 |
| TH28051 | *flhB*$_{\Delta CCD}$ | 1015 | 18 | 1.8 |
| TH28054 | Δ*flk flhB*$_{\Delta CCD}$ | 654 | 12 | 1.8 |
| TH29314 | Δ*flk flhB*N269A | 1305 | 20 | 1.5 |
| TH28053 | Δ*flgHI* Δ*flk* | 1405 | 288 | 20.5 |
| TH28055 | Δ*flgHI flhB*$_{\Delta CCD}$ | 954 | 3 | 0.3 |
| TH29315 | Δ*flgHI flhB*N269A | 1560 | 0 | <0.01 |
| TH28056 | Δ*flgHI* Δ*flk flhB*$_{\Delta CCD}$ | 704 | 14 | 2 |
| TH29316 | Δ*flgHI* Δ*flk flhB*N269A | 998 | 2 | 0.2 |
| TH29317 | *flhD*C* flhB*$_{\Delta CCD}$ Δ*flk* | 946 | 764 | 90 |
| TH29318 | *flhD*C* flhB*$_{\Delta CCD}$ Δ*flgHI* | 1180 | 1 | 0.1 |
| TH29319 | *flhD*C* flhB*$_{\Delta CCD}$ Δ*flk* Δ*flgHI* | 1480 | 1465 | 99 |
| TH29357 | *flhD*C** Δ*flhB* Δ*flk* | 950 | 0 | <0.01 |
| TH29358 | *flhD*C** Δ*flhB* Δ*flgHI* | 1000 | 0 | <0.01 |
| TH29359 | *flhD*C** Δ*flhB* Δ*flk* Δ*flgHI* | 1200 | 0 | <0.01 |

*Cells were imaged at cell density of approximately 1.

**Table S4. Strains used in Figure 7 and supplemental Figure 5:**

**<u>FlgM-Bla</u> strains:**
All strains contain *flgM6407*::*bla* Δ*flgB-L8735 fliA5225* Δ*fliB-T7771 fljB*$^{enx}$ *vh2*

| | WT | FliP (I95N) | FliQ (G32D) | FliR (Q210 stop) | FliR (V215D) | FliR (T221DUP) |
|---|---|---|---|---|---|---|
| *flk*$^{+}$ | TH29405 | TH29427 | TH29431 | TH29439 | TH29443 | TH29447 |
| Δ*flk* | TH29406 | TH29428 | TH29432 | TH29440 | TH29444 | TH29448 |
| Δ*fliK* | TH29684 | TH29451 | TH29455 | TH29463 | TH29467 | TH29471 |
| Δ*fliK* Δ*flk* | TH29685 | TH29452 | TH29456 | TH29464 | TH29468 | TH29472 |

**<u>FlgE-Bla</u> strains:**
All strains contain *flgE6569*::*bla* Δ*flgBC6557 fliA5225* Δ*fliB-T7771 fljB*$^{enx}$ *vh2*

| | WT | FliP (I95N) | FliQ (G32D) | FliR (Q210 stop) | FliR (V215D) | FliR (T221DUP) |
|---|---|---|---|---|---|---|
| *flk*$^{+}$ | TH29407 | TH29429 | TH29433 | TH29441 | TH29445 | TH29449 |
| Δ*flk* | TH29408 | TH29430 | TH29434 | TH29442 | TH29446 | TH29450 |
| Δ*fliK* | TH29686 | TH29453 | TH29457 | TH29465 | TH29469 | TH29473 |
| Δ*fliK* Δ*flk* | TH29687 | TH29454 | TH29458 | TH29466 | TH29470 | TH29474 |

**<u>FliK-Bla</u> strains:**
All strains contain *fliK7582*::*bla flhB7152*(N269A) Δ*flgB-L8735 fliA5225* Δ*fliB-T7771 flhD8070 flhC8092 fljB*$^{enx}$ *vh2*

| | WT | FliP I95N | FliQ (G32D) | FliR Q210 stop | FliR T221DUP | FliR V215D | FliK P296L | FliK Δ311-320 |
|---|---|---|---|---|---|---|---|---|
| *flk*$^{+}$ | TH30353 | TH30354 | TH30355 | TH30357 | TH30359 | TH30358 | TH30405 | TH30403 |
| Δ*flk* | TH30360 | TH30361 | TH30362 | TH30364 | TH30366 | TH30365 | TH30406 | TH30404 |
